# Cerebellar lesions disrupt spatial and temporal visual attention

**DOI:** 10.1101/822635

**Authors:** B.T. Craig, A. Morrill, B. Anderson, J. Danckert, C.L. Striemer

## Abstract

The current study represents the first comprehensive examination of spatial, temporal and sustained attention following cerebellar damage. Results indicated that, compared to controls, cerebellar damage resulted in a larger cueing effect at the longest SOA – possibly reflecting a slowed the onset of inhibition of return (IOR) during a reflexive covert attention task, and reduced the ability to detect successive targets during an attentional blink task. However, there was little evidence to support the notion that cerebellar damage disrupted voluntary covert attention or the sustained attention to response task (SART). Lesion overlay data and supplementary voxel-based lesion symptom mapping (VLSM) analyses indicated that impaired performance on the reflexive covert attention and attentional blink tasks were related to damage to Crus II of the left posterior cerebellum. In addition, subsequent analyses indicated our results are not due to either general motor impairments or to damage to the deep cerebellar nuclei. Collectively these data demonstrate, for the first time, that the same cerebellar regions may be involved in both spatial and temporal visual attention.

---

Traditionally, the cerebellum is considered to be important for the timing and coordination of motor outputs and motor learning (Glickstein, Strata, & Voogd, 2009; Glickstein, Sultan, & Voogd, 2011). However, more recent research has highlighted a role in a diverse array of cognitive, affective, and perceptual processes, including language, working memory, executive control, emotion and motion perception (Adamaszek et al., 2017; O. Baumann et al., 2015; Bellebaum & Daum, 2007; Marvel & Desmond, 2010; Sacchetti, Scelfo, & Strata, 2009; Schmahmann, Guell, Stoodley, & Halko, 2019; Schmahmann & Sherman, 1998; Stoodley, MacMore, Makris, Sherman, & Schmahmann, 2016; Stoodley & Schmahmann, 2009a; Stoodley & Stein, 2011).

The role of the cerebellum in cognition is supported by anatomical evidence demonstrating connections from the ventral dentate nucleus to non-motor regions of posterior parietal and prefrontal cortex (Clower, West, Lynch, & Strick, 2001; Dum & Strick, 2003; Strick, Dum, & Fiez, 2009). In addition, studies examining functional connectivity have revealed a number of distinct networks in the cerebellum that are functionally connected to different cortical networks known to be involved in a variety of cognitive functions (Buckner, Krienen, Castellanos, Diaz, & Yeo, 2011; Wang, Buckner, & Liu, 2013).

One area of contention regarding the cerebellum’s role in cognition concerns its potential involvement in attention. Courchesne and colleagues first demonstrated that patients with cerebellar lesions were slower to shift attention between streams of auditory or visual events (Akshoomoff & Courchesne, 1992, 1994) and were slower to orient attention towards peripheral targets (Townsend et al., 1999). Subsequent studies, however, failed to identify clear attentional deficits in cerebellar patients (Dimitrov et al., 1996; Golla, Thier, & Haarmeier, 2005; Yamaguchi, Tsuchiya, & Kobayashi, 1998). This has led some to suggest that initial findings likely reflected slowed motor responses (i.e., slowed button presses or eye movements) masquerading as attentional impairments (Glickstein et al., 2011; Haarmeier & Thier, 2007; Ravizza & Ivry, 2001).

There are a number of potential reasons for these conflicting findings. First and foremost, a majority of studies have included a mix of patients with cerebellar lesions, degeneration, and development disorders (Glickstein et al., 2011; Haarmeier & Thier, 2007; Ravizza & Ivry, 2001). Such an approach implicitly assumes that all regions of the cerebellum are *equally involved* in all aspects of the cognitive task being tested. That is, lesion location was, at least initially, not given serious consideration in the analysis or the model of cerebellar involvement in cognition. More recent neuroimaging research, and studies examining patients with circumscribed cerebellar lesions, suggest that distinct cerebellar regions are involved in spatial and non-spatial attention (Baier, Dieterich, Stoeter, Birklein, & Muller, 2010; Schweizer, Alexander, Cusimano, & Stuss, 2007; Striemer, Cantelmi, Cusimano, Danckert, & Schweizer, 2015; Striemer, Chouinard, Goodale, & de Ribaupierre, 2015; Townsend et al., 1999).

Specifically with respect to visuospatial attention, a number of studies have employed the well-known covert attention paradigm developed by Posner and colleagues (Posner, Rafal, Choate, & Vaughan, 1985; Posner, Snyder, & Davidson, 1980). In this task, participants fixate centrally while detecting peripheral targets that can appear at a previously cued location (i.e., a valid trial) or in the location opposite the cue (i.e., an invalid trial). Cues can be either predictive of the impending target location (presumably evincing voluntary allocation of attention) or non-predictive (evincing a reflexive orienting processes). Previous studies showed deficits in covert attention for predictive cues (Baier et al., 2010; Townsend et al., 1999). However, a recent patient study from our group (Striemer, Cantelmi, et al., 2015) demonstrated that patients with lateral cerebellar lesions showed deficits of reflexive covert orienting. Specifically, cerebellar patients showed smaller cueing benefits at short SOAs (50ms) and a trend towards a diminished inhibition of return (IOR) – the typical reversal of the cueing effect found at a longer SOAs (Klein, 2000; Posner et al., 1985).

Functional neuroimaging work from our group supports the role of lateral cerebellar regions in reflexive covert orienting (Striemer, Chouinard, et al., 2015). We found significant BOLD activation in lobule VI of the left cerebellum for both reflexive and voluntary covert attention, with or without eye movements and controlling for manual responses. Importantly, activation in the cerebellar ROI was greater for reflexive compared to voluntary attention, and was significantly correlated with increased BOLD activity in superior and inferior parietal lobes, and the frontal eye fields – nodes of the fronto-parietal attention network.

Spatial orienting represents just one kind of attentional process. Attention must also be oriented in time and sustained over time (Husain & Nachev, 2007; Robertson, Manly, Andrade, Baddeley, & Yiend, 1997). Previous research demonstrated an impairment in performance on the attention blink (AB) task – a widely utilized measure of the temporal allocation of attention (Dux & Marois, 2009) – following cerebellar damage. Here participants must identify two targets within a rapid stream of stimuli presented in central vision (Dux & Marois, 2009; Raymond, Shapiro, & Arnell, 1992). Schweizer and colleagues (2007) found that cerebellar patients had a larger AB. That is, they were less accurate at detecting the second target when it appeared shortly after the first. Importantly, patients achieved equivalent target two accuracy levels at about the same time as healthy controls. In other words, their AB was larger in magnitude, but was not prolonged in duration as it is for patients with inferior parietal or superior temporal damage (Husain, Shapiro, Martin, & Kennard, 1997; Shapiro, Hillstrom, & Husain, 2002).

To our knowledge, only one single case study has examined sustained attention following cerebellar damage (Schweizer et al., 2008). Fundamentally, sustained attention tasks require prolonged focus of attention over longer periods of time, without necessarily taxing the temporal allocation of attention itself (Esterman & Rothlein, 2019). Previous research demonstrated that the cerebellum is involved in executive functions (Bellebaum & Daum, 2007), as well as sequence detection (Leggio & Molinari, 2015; Molinari et al., 2008) and performance monitoring (Peterburs & Desmond, 2016). Thus, it is plausible that cerebellar damage could disrupt the ability to maintain performance on a task over time, which is a core component of sustained attention.

Here we contrasted performance on three standard attention tasks in patients with cerebellar damage and age-matched controls. Participants completed two versions of the Posner cueing paradigm to examine reflexive and voluntary orienting with a view to replicating and extending our prior work. Next, we examined the temporal allocation of attention and sustained non-spatial attention by having participants complete versions of the AB task and the Sustained Attention to Response (SART) task (Manly, Robertson, Galloway, & Hawkins, 1999; Robertson et al., 1997). Lesion overlay data (and supplementary lesion analyses) were used to identify specific regions of the cerebellum linked to observed deficits. The current study is the first comprehensive investigation of the effects of cerebellar lesions on spatial and non-spatial attention in the same patient group. Our results demonstrate, for the first time, that lesions to Crus II of the left cerebellum disrupt reflexive spatial attention and the temporal allocation of attention (the AB); However, there was less support for an influence of cerebellar lesions on voluntary spatial attention or sustained attention.

## Methods

### Participants

Fourteen patients with cerebellar lesions participated in the current study (mean age 63.57 years; SD=12.57; range 38-83; 6 females). All lesions were a result of a single isolated cerebellar stroke classified as either a left (n=8), right (n=2), or bilateral (n=4) based on clinical notes and confirmed via MRI (n=8) or CT scans (n=6). All patients were right-handed and had no additional neurological deficits. Testing occurred in the chronic stages post-stroke (mean time post stroke= 4.5 years; Table 1). Patients were recruited form the Neurological Patient Database maintained by the University of Waterloo (Heart and Stroke Foundation funded) in which the patients had previously provided consent to be contacted for research studies. It is important to note that, although this is not a very large patient sample, isolated cerebellar strokes comprise less than 3% of all stroke cases making it difficult to recruit large number of these patients (Kelly et al., 2001; Macdonell, Kalnins, & Donnan, 1987; Tohgi, Takahashi, Chiba, & Hirata, 1993).

**Table 1:**
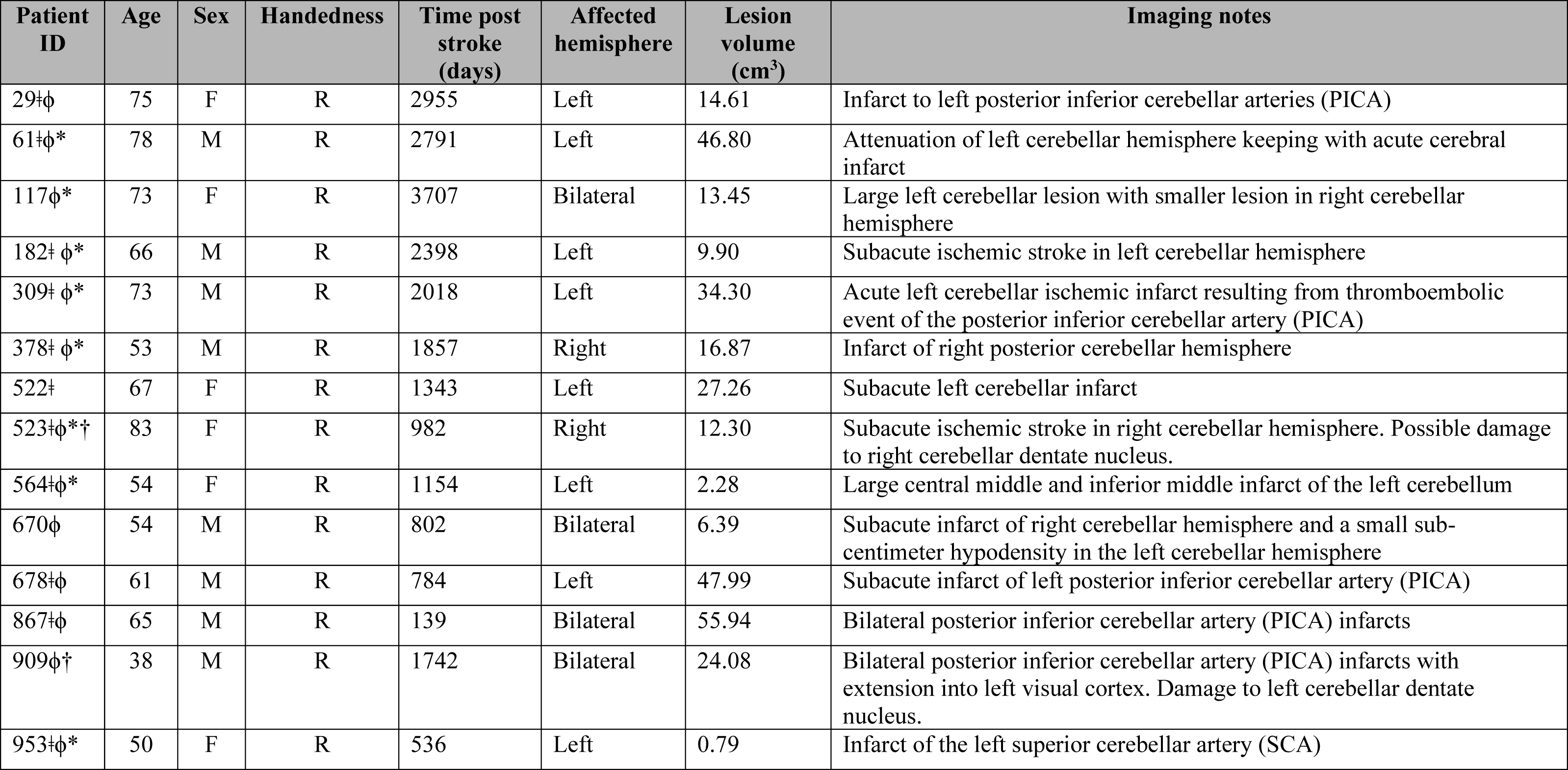
Clinical data for the 14 cerebellar patients. Symbols: ǂ completed the covert attention tasks; ϕ completed the non-spatial attention tasks; * indicates that a new high-resolution MRI was acquired. † indicates damage to the cerebellar dentate nucleus.

| Patient ID | Age | Sex | Handedness | Time post stroke (days) | Affected hemisphere | Lesion volume (cm<sup>3</sup>) | Imaging notes |
| --- | --- | --- | --- | --- | --- | --- | --- |
| 29‡ϕ | 75 | F | R | 2955 | Left | 14.61 | Infarct to left posterior inferior cerebellar arteries (PICA) |
| 61‡ϕ* | 78 | M | R | 2791 | Left | 46.80 | Attenuation of left cerebellar hemisphere keeping with acute cerebral infarct |
| 117ϕ* | 73 | F | R | 3707 | Bilateral | 13.45 | Large left cerebellar lesion with smaller lesion in right cerebellar hemisphere |
| 182‡ϕ* | 66 | M | R | 2398 | Left | 9.90 | Subacute ischemic stroke in left cerebellar hemisphere |
| 309‡ϕ* | 73 | M | R | 2018 | Left | 34.30 | Acute left cerebellar ischemic infarct resulting from thromboembolic event of the posterior inferior cerebellar artery (PICA) |
| 378‡ϕ* | 53 | M | R | 1857 | Right | 16.87 | Infarct of right posterior cerebellar hemisphere |
| 522‡ | 67 | F | R | 1343 | Left | 27.26 | Subacute left cerebellar infarct |
| 523‡ϕ*† | 83 | F | R | 982 | Right | 12.30 | Subacute ischemic stroke in right cerebellar hemisphere. Possible damage to right cerebellar dentate nucleus. |
| 564‡ϕ* | 54 | F | R | 1154 | Left | 2.28 | Large central middle and inferior middle infarct of the left cerebellum |
| 670ϕ | 54 | M | R | 802 | Bilateral | 6.39 | Subacute infarct of right cerebellar hemisphere and a small sub-centimeter hypodensity in the left cerebellar hemisphere |
| 678‡ϕ | 61 | M | R | 784 | Left | 47.99 | Subacute infarct of left posterior inferior cerebellar artery (PICA) |
| 867‡ϕ | 65 | M | R | 139 | Bilateral | 55.94 | Bilateral posterior inferior cerebellar artery (PICA) infarcts |
| 909ϕ† | 38 | M | R | 1742 | Bilateral | 24.08 | Bilateral posterior inferior cerebellar artery (PICA) infarcts with extension into left visual cortex. Damage to left cerebellar dentate nucleus. |
| 953‡ϕ* | 50 | F | R | 536 | Left | 0.79 | Infarct of the left superior cerebellar artery (SCA) |

For comparison, we tested a total of 24 age-appropriate healthy controls (mean age=70.95 years; SD=8.92; range 53-83; 19 females). The two groups did not differ in mean age (t(21)=1.93, *p*=.07). Control participants were all right-handed and had no history of neurological impairment. Healthy age-appropriate controls were recruited from either the Waterloo Research in Aging Pool (WRAP; Waterloo, Ontario, Canada), or from the MINERVA Senior Studies Institute at MacEwan University (Edmonton, Alberta, Canada). All participants gave written consent prior to the first testing session.

This project was approved by the MacEwan University Research Ethics Board, the University of Waterloo Research Ethics Board, and the Tri-Hospital Research Ethics Board (Kitchener-Waterloo, Ontario). All participants were compensated $10 per hour for each session.

### General procedures

All participants completed four attention tests: reflexive and voluntary covert attention, the AB, and the SART. Each test is described in detail below. All participants completed the four tasks over two separate testing sessions. All button-press responses (when required) were made with the right (dominant) hand. One session tested covert spatial attention (reflexive followed by voluntary covert attention) and the other session tested non-spatial attention (AB and SART; counterbalanced). We always tested reflexive prior to voluntary covert attention in order to avoid any potential carryover effects of the predictive cue contingency in the voluntary task from influencing performance on the reflexive task. The order of the testing sessions (spatial vs. non-spatial attention) was counterbalanced between patients and controls. Some patients (n=8) attended a third session where we obtained a high-resolution T1 MRI anatomical scan of their brain. The remaining patients (n=6) were either unable to attend a third session or were precluded from having an MRI for medical reasons. To assess the presence of motor deficits in the cerebellar patients we administered a modified version of the International Cooperative Ataxia Rating Scale (ICARS Trouillas et al., 1997). The ICARS assessment was administered in the same session as the covert attention tasks in order to keep the two sessions roughly the same length.

### International cooperative ataxia rating scale (ICARS)

The ICARS (Trouillas et al., 1997) examines a patient’s walking capacity, gait speed, standing capacities and balance. The test also examines dysmetria and intention tremor in each of the upper and lower limbs using the finger-to-nose test, as well as the heel-to-toe test and the timing and coordination of limb movements using alternating pronation and supination of the hands. Finally, the ICARS assesses oculomotor functions by searching for evidence of gaze-evoked nystagmus, deficits in oculomotor pursuit, or saccadic dysmetria. All tests were scored using the established ICARS scoring procedure (Trouillas et al., 1997). The modified ICARS had a total possible score of 56 (18 (posture and gait) + 32 (limb coordination) + 6 (oculomotor functions)) with higher scores indicative of greater impairment. All ICARS assessments were video recorded for offline analysis and confirmed by two separate raters to check for inter-rater reliability.

### Apparatus

All attention tasks were administered in a dimly lit room on a PC laptop computer with a 53cm x 30cm screen (1920 x 1080 resolution, 60 Hz refresh rate), while resting their head in a chin rest placed 57cm from the screen. All tasks were run using Superlab 5 software (https://www.cedrus.com/superlab/) with responses collected using a Cedrus RB-730 response pad with ± 2-3ms reaction time resolution.

### Reflexive covert attention

For the reflexive covert attention task (Figure 1) a 1cm x 1cm white fixation cross was presented centrally on a uniform black background. Two white boxes 2cm x 2cm size, located 10cm (10°) to the left and right, represented potential target locations. Box size was increased to 2.5cm x 2.5 cm to function as a cue. Targets were an asterisk (‘*’) 1 cm in diameter presented in the center of one of the two boxes. Participants were asked to fixate centrally while attending to the boxes to the left and right.

**Figure 1.**
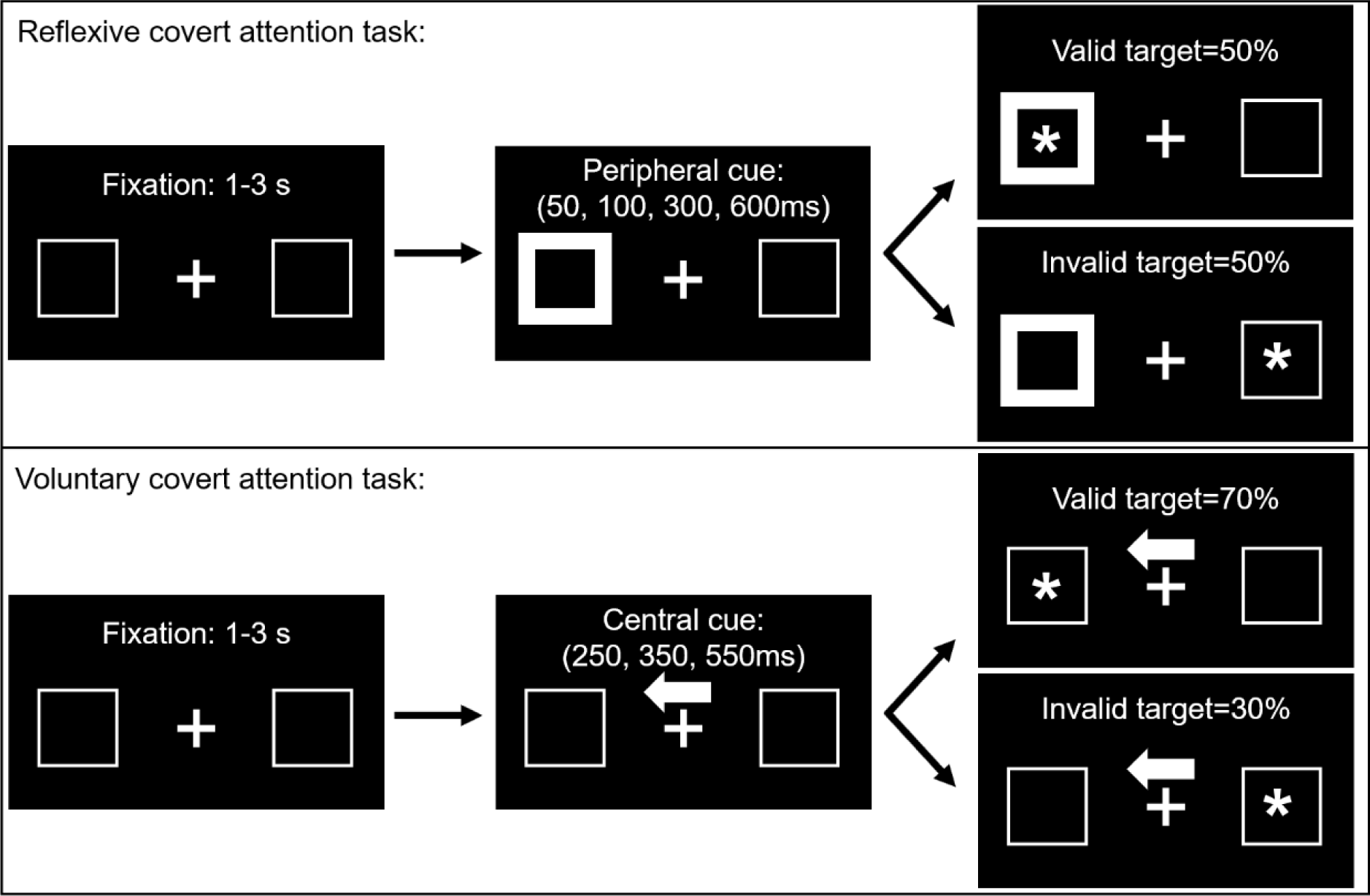
For the reflexive covert attention task (top panel) a single trial began with a fixation period of 1-3s followed by a peripheral cue on the left or right. Following a stimulus onset asynchrony (SOA) of 50, 100, 300 or 600ms a target (*) subsequently appeared either at the cued (i.e., valid) or uncued (i.e., invalid) location with equal probability. For the voluntary covert attention task (bottom panel) a central arrow cue was used, and the target appeared at the cued location on 70% of trials following an SOA of either 250, 350 or 550ms. Note that for both tasks the cue remained on the screen until the target appeared.

Fixation was monitored (and recorded) using a Logitech 720 HD webcam zoomed in on the participant’s eyes. In addition, participants were periodically reminded to maintain fixation. We discarded a trial due to an eye movement if an eye movement that was large enough to bring the target into central vision was initiated during the time between when the target appeared, and the response was made. Prior to covert attention testing sessions, we asked each patient to fixate centrally and then to make saccades between the fixation point and the target locations so we would know how large the saccade would need to be in order to bring the target into central vision. Given that targets were located 10° to the left and right, any eye movements toward a target were easily detectable on the webcam. Eye movements were analyzed offline by two independent raters to establish inter-rater reliability.

Each trial began with a 1000 Hz tone followed by a fixation period of between 1 to 3 seconds (randomly selected equally often at 500ms increments). Following fixation, one of the peripheral boxes appeared to brighten acting as a reflexive cue to attract attention to that location. Following a stimulus-onset-asynchrony (SOA) of either 50, 100, 300, or 600 ms the target appeared at either the cued location (i.e., “valid trials”), or the uncued location (i.e., “invalid trials”). The peripheral cue remained present until the target appeared and was not predictive of the target’s location (i.e., 50% valid). Participants responded via button press as quickly and accurately as possible following target onset. In a previous study (Striemer, Cantelmi, et al., 2015) we observed a trend towards a reduced IOR at a 300ms SOA. In the current study we added a 600ms SOA to examine whether this trend would become significant.

Participants completed two blocks of 190 trials, which consisted of 10 validly and invalidly cued left and right targets for each of the four SOAs, as well as 10 trials in which no cue was presented prior to target onset (i.e., ‘no cue’ trials). Each block also contained 10 ‘catch’ trials in which the cue was presented but no target appeared. Catch trials were used to ensure that participants were reacting to the onset of the target and not the cue. Participants responded on 3% (or less) of catch trials. Participants completed 12 practice trials prior to completing the main experiment to ensure they understood the task.

### Voluntary covert attention (spatial attention)

The setup for the voluntary covert attention task (Figure 1) was similar to the reflexive covert attention task, with a few differences. First, target locations were cued via a central arrow symbol (1cm tall x 2cm wide) that accurately predicted the target location on 70% of trials. This version of the covert attention task utilized SOAs of 250, 350, and 550 milliseconds. Participants completed two blocks of 130 trials with 12 invalid trials and 28 valid trials for each SOA for both left and right targets. In addition, we also included 10 no cue trials and 10 ‘catch’ trials. Participants were specifically told about the predictive nature of the cue. Although we had no a priori hypotheses regarding SOA for the voluntary covert attention task, we nevertheless included three SOAs in order to be able to plot trends over time should they exist.

For both covert attention tasks the primary dependent measure was the reaction time (RT) to respond to each trial type (valid vs. invalid) by SOA combination. In addition, to analyze the effect of cue validity as a function of SOA, we calculated cue effect sizes (CES) for each cue x SOA combination by subtracting the RT for valid trials from the RT for invalid trials. The CES represents the benefit of the cue while controlling for overall response speed such that a positive CES reflect a cueing benefit (i.e., faster RTs) for validly cued trials, whereas a negative CES reflects a cueing benefit for invalidly cued trials (i.e., IOR). Reaction times that were below 150ms or more than 2 SD above or below the participant’s mean for that cue-target-SOA combination were considered outliers and were removed from the analysis. For controls, any participant whose CES was more than 2SD above or below the mean for two or more SOAs was removed from the dataset as an outlier. This was determined prior to data analysis.

### Attentional blink (AB) task

For the attentional blink (AB) task (Figure 2) we adopted the procedure utilized by Schweizer and colleagues (2007). Participants were presented with an RSVP stream of individual uppercase white or red letters on a black background (67-point font) at a rate of 130ms per letter. Within the AB task there were two types of trials, 1-Target trials and 2-Target trials. 2-Target trials were essential for eliciting the AB effect, whereas 1-Target trials were used as a control condition.

**Figure 2.**
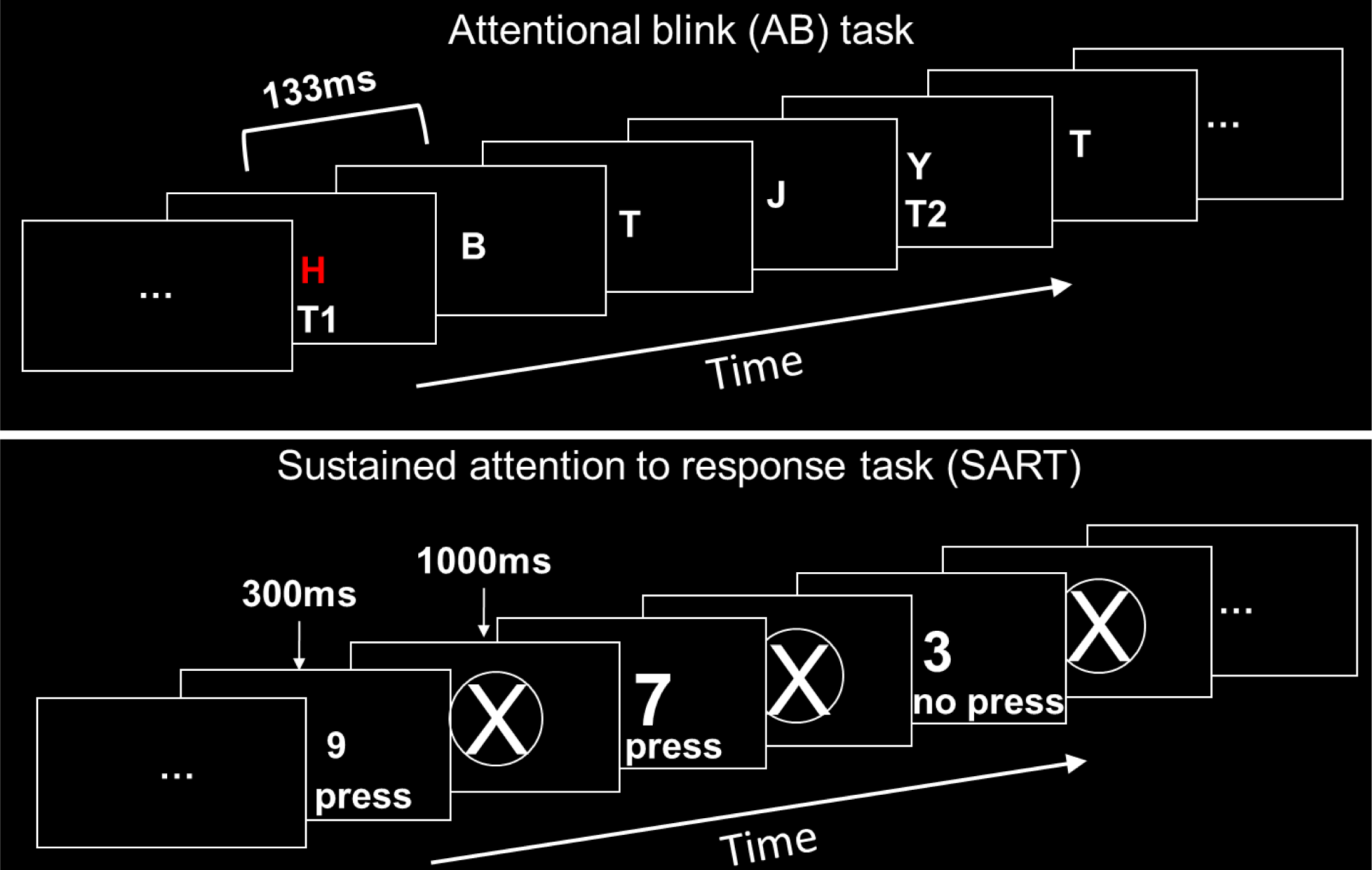
During the attentional blink (AB; top panel) task participants were presented with a rapid serial visual presentation (RSVP; 133ms/letter) where they were asked to indicate the presence of a red H or S or a white X or Y. On 1-Target trials only a white X or Y was presented. On 2-Target trials a white X or Y was presented either 1, 2, 3, 4, 8, or 12 letters after the presentation of a red H or S. For the sustained attention to response task (SART; bottom panel) participants were to press a button each time a number appeared on the screen except for ‘3’. Each number appeared for 300ms followed by a mask period of 1000ms.

For all 2-Target trials participants were instructed to report the presence of target letters which would always consist of a red H or S (T1) followed by a white X or Y (T2). T1 (red H or S) always appeared prior to T2 (white X or Y) in the letter stream. Non-target letters were drawn from the remaining letters of the alphabet (randomly selected) and were presented in white. In a single 2-Target trial 6-9 non-target letters (with equal probability) were presented prior to T1 (a red H or S), with equal probability. Then a total of 9-12 letters were presented (with equal probability) following T1 in which T2 (a white X or Y, with equal probability) could appear at each of 6 positions or ‘lags’ (1, 2, 3, 4, 8, or 12 letters after T1, with equal probability). In addition, there were always 1-4 letters presented after T2. Following the letter stream participants were asked to indicate whether any red target letters were presented (H or S, or “no”), and whether a white X or Y was presented. After each trial the researcher coded the participant’s responses using the keyboard before pressing the spacebar to move to the next trial. For 2-Target trials the primary dependent measure was the participant’s accuracy in identifying T2 after correctly identifying T1 as a function of lag.

1-Target trials were similar to 2-Target trials except that, for 1-Target trials, T1 (the red H or S) was replaced by a white non-target letter (randomly selected) which was then followed by T2 (a white X or Y) presented under the same constraints as in 2-Target trials (described above). Following the letter stream participants were again asked to identify if any red letters were presented (H, S or “no”) and whether a white X or Y was presented. The primary dependent measure on these trials was the accuracy in identifying T2 in the absence of T1.

Participants performed 180 trials with 1/3 being 1-Target trials, and 2/3 being 2-Target trials. 1-Target and 2-Target trials were intermixed and randomly presented. Participants were given 10 practice trials before completing the main task and were offered a short break halfway through. Control participants were removed as outliers if their accuracy for either the 1-Target or 2-Target tasks was more than 2SD outside of the overall group mean for a given Lag. The primary dependent measures for the AB task were accuracy in detecting a single target on 1-Target trials and accuracy in detecting both T1 and T2 on 2-Target trials. For controls, if their accuracy for 2 or more lags was more than 2SD above or below the group mean for that lag, they were removed as an outlier. This was determined prior to data analysis.

### Sustained attention to response task (SART)

In the SART the digits 1-9 were presented one at a time in the center of the screen in random order. Each digit was presented for 300ms, followed by a 1000ms mask. Participants pressed the space bar of the computer keyboard as quickly and accurately as possible for every number except ‘3’. When a ‘3’ appeared, participants were told to withhold their response. All digits were a standard white Arial font that varied in size (48, 72, 94, 100 or 120pt font) and were presented on a black background. The mask was a large white circle with an X in the center. There were 225 numbers presented in total with each number presented 25 times in random order (25 of them being 3). Participants were given 15 practice trials prior to the experimental trials to ensure they understood the task. Here, we measured the percentage of commission errors (i.e., presses for ‘3’), and misses (or omission errors – failing to press for numbers other than 3), as well as the RTs for errors and correct responses. Responses that were under 150ms were discarded as anticipatory. For controls, any participant whose errors of commission (presses for 3) were more than 2SD above the control mean were removed as an outlier. This was determined prior to data analysis.

### Lesion data

To explore the association between lesion location and behaviour we acquired medical imaging data for each of the 14 patients and created a lesion overlay plot. For 8 patients we acquired high resolution 160 slice 1mm ISO-voxel T1-weighted MRI scans collected on a 1.5T Philips scanner (Grand River Hospital, Kitchener, Ontario). The remaining six patients were either unable to undergo an MRI scan due to safety concerns (e.g., surgical implants, metal in their body), or elected not to do so for personal reasons. For these patients we acquired existing clinical MRI or CT scan data from their medical records. For all patients, lesions were traced by an experienced neurologist (B.A.) using MRIcron software (http://people.cas.sc.edu/rorden/mricron/index.html). All patient anatomicals and lesion masks were normalized into MNI space onto a high-resolution CT template using the Clinical Toolbox for SPM 12 (Rorden, Bonilha, Fridriksson, Bender, & Karnath, 2012). The normalized individual lesion masks were then combined to make a group lesion mask in MRIcron which was overlaid onto the same high-resolution CT template. We then extracted the MNI coordinates for the regions where the largest number of patients had overlapping lesions and converted them into Talairach coordinates. These Talairach coordinates were used to localize the lesioned regions using the Talairach Daemon Atlas (http://www.talairach.org/). In addition to examining lesion overlap for the overall group, we also examined each patient’s scan for damage to the cerebellar dentate output nuclei using the probabilistic dentate atlas developed by Dimitrova and colleagues (Dimitrova et al., 2006).

Finally, in an attempt to link patients’ behavioural performance with specific cerebellar regions we used voxel-based lesion symptom mapping (VLSM) – a technique that allows researchers to examine the relationship between damage to specific voxels in the brain with behavior (Bates et al., 2003). To do this we used the NPM toolbox that is included as part of MRIcron (Rorden, Karnath, & Bonilha, 2007). Although these analyses are admittedly underpowered with our patient sample size, we have included them in the Supplementary Material. Importantly, the VLSM data largely confirm the results of our lesion overlay plots.

### Statistical analyses

Statistical analyses were carried out using JASP (JASP-Team, 2020). All within-subject ANOVAs were computed using a Greenhouse-Geisser correction when necessary (Greenhouse & Geisser, 1959) and all post-hoc tests were carried out using the Holm procedure (Holm, 1979) to control for familywise error rate (*p*<.05). Finally, for effects of interest we also report the results of Bayesian t-tests to better understand the strength of evidence our data provides in support of the alternative or null hypothesis (Jarosz & Wiley, 2014; Masson, 2011; Wagenmakers, 2007). We report inverse Bayes factors (BF_10_) in which larger numbers (i.e., >1) indicate more support for the alternative hypothesis and small numbers (i.e., <1) indicate more support for the null hypothesis. For example, a BF_10_= 5.00 would mean that the alternative hypothesis is five times more likely than the null hypothesis. The details of these analyses are reported in the Supplementary Material.

No part of the procedures or analysis for this study was pre-registered prior to the research being conducted. All of the manipulations and measures collected in this study are reported in this manuscript.

### Data availability statement

The authors confirm that the summarized group data supporting the findings of this study are available within the article and its Supplementary Material. Raw data and individual participant data cannot be made available because of ethical restrictions. Specifically, all participants in the study signed a consent form which explicitly indicated that, “only the researchers involved in the study will have access to this data.” Requests for access to individual participant data must be submitted to the corresponding author, and a data sharing agreement must be submitted to the University of Waterloo Office of Research Ethics.

### Digital study materials

All experimental stimuli and presentation files have been archived on <u>Open Science Framework (OSF)</u>.

## Results

### International Cooperative Ataxia Rating Scale (ICARS)

We acquired International Cooperative Ataxia Rating Scale (ICARS) scores for 12 of the 14 patients, with one patient dropping out of the study prior to completing this portion and data for a second patient lost due to experimenter error. ICARS data are shown in Table 2. There was high inter-rater reliability for overall ratings on the ICARS (r=.98, *p*<.0001).

**Table 2.**
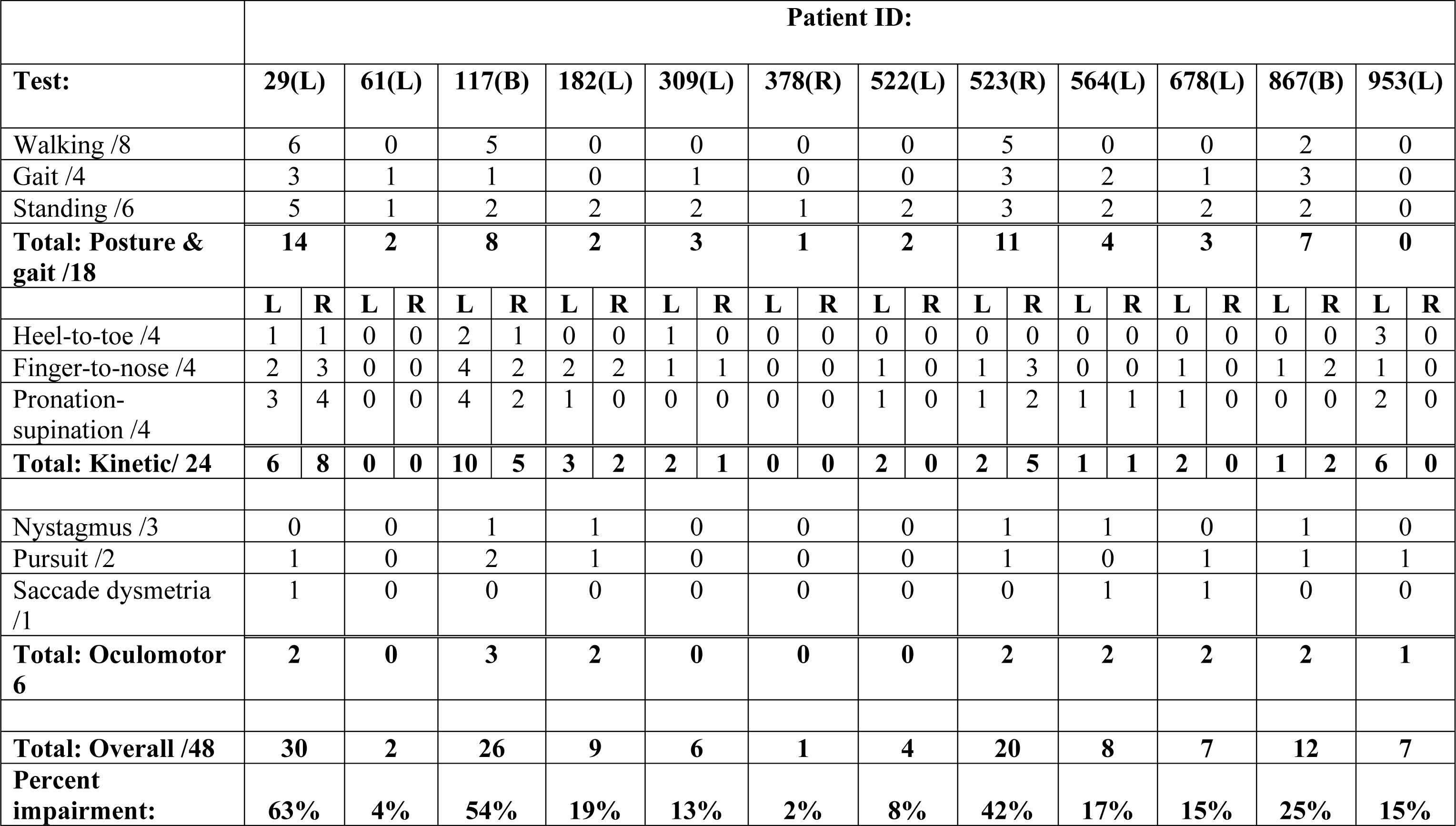
International cooperative ataxia rating scale (ICARS) data for the cerebellar patients (n=12). Total scores for each subtest and overall scores are listed for each patient (side of lesion in brackets). Larger numbers indicate greater motor impairment. Percentage impairment scores were calculated by dividing the patient’s raw score by total possible score (Trouillas et al., 1997).

### Spatial attention tasks

All RT data for the covert attention tasks for each patient and the controls are presented in Supplementary Tables 1 & 2. Of the 14 patients recruited, data was collected for the covert spatial attention tasks on only 11 as the remaining three patients had lesions extending into occipital cortex resulting in partial peripheral vision loss that made it difficult to view one of the peripheral boxes when fixating. Which patients completed each attention task is noted in Table 1. One control participant had a CES that was more than 2SD above the control mean in two different SOAs for both the reflexive and voluntary attention tasks and was removed from both datasets as an outlier (see Methods) resulting in a total of 23 controls in the final analysis.

Overall, participants did an excellent job fixating with fewer than 8% of trials excluded due to fixation problems. There was high inter-rater reliability for saccades (r=.89, *p*<.001).

### No-cue trials

Prior to analyzing the data from the cued trials in the covert attention tasks we first analyzed the data from the no-cue trials. To analyze these data we used a mixed-model ANOVA with side of target (left vs. right) and task (reflexive vs. voluntary) as the within-subject factors and group (patients vs. controls) as a between-subject factor. This analysis revealed a main effect of side of target (F(1,32)=23.90, *p*<.001, η^2^_p_= .43) such that left sided targets (555ms) were responded to more quickly than right sided targets (616ms). No other main effects or interactions with group were significant (all *p*’s >.11). Given that there were no interactions with group all subsequent analyses were collapsed across side of target.

### Reflexive covert attention

A mixed-model ANOVA with group (patients vs. controls) as a between-subject factor, and cue (valid vs. invalid), and SOA (50, 100, 300, 600) as within-subject factors, revealed main effects of cue (F(1,32)=97.03, *p*<.001, η^2^_p_=.75), SOA (F(2.06,65.89)=4.62, *p*=.021, η^2^_p_=.11), and a marginal main effect of group (F(1,32)=4.01, *p*=.054, η^2^_p_ =.11). Specifically, RTs were faster for valid (542ms) compared to invalid (584ms) trials, RTs were slower overall at the 50ms SOA (574ms) compared to the 300 (556ms) and 600ms SOAs (557ms; all *p’s* <.024, Holm-corrected), and RTs were somewhat slower for patients (599ms) compared to controls (526ms).

There was also a significant cue x SOA x group interaction (F(3,96)=4.20, *p*=.008, η^2^_p_ =.12; Figure 3A). A simple-main effects analysis revealed that this interaction was driven by a difference in CES at 600 ms SOA. Patients showed a large, positive CES (mean=54ms) compared to controls (mean=-3 ms; t(32)=3.48, *p*=.004, d=1.28, Holm-corrected). Furthermore, whereas the CES at the 600ms SOA was significantly larger than zero for patients (t(10)=2.50, *p*=.031, d=.76) the CES for controls was not different from zero (t(22)=.54, *p*=.60, d=.11). CES sizes at all other SOAs did not differ between groups (all *p*’s >.14, uncorrected; Figure 3B). These results were supported by Bayesian independent samples t-tests which indicated strong support for a difference in CES at the 600ms SOA (BF_10_=22.49) but anecdotal support for the null hypothesis for the other SOAs (all BF_10_= <.80; see Supplementary Material).

**Figure 3.**
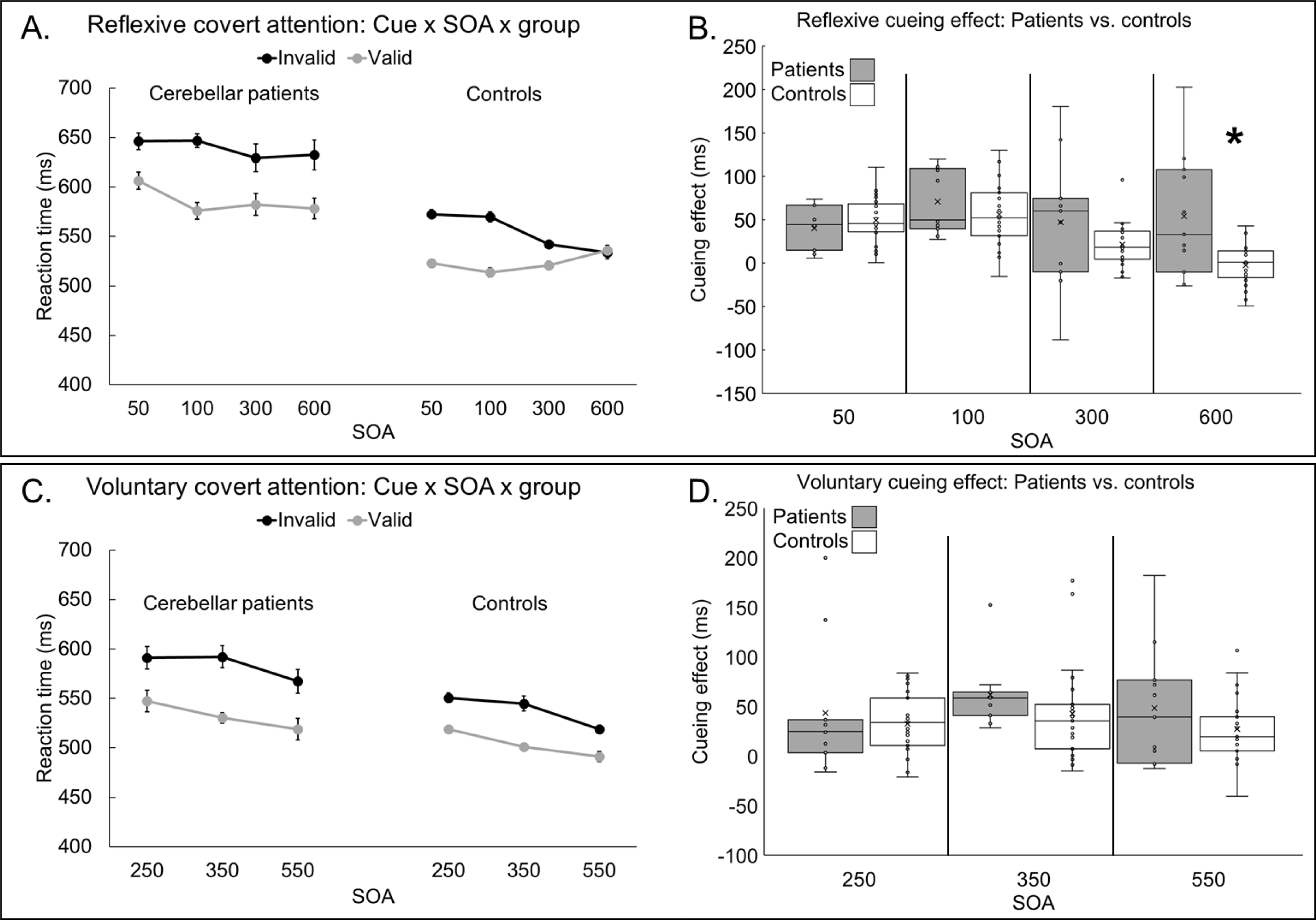
**Reaction time (RT) data** for the reflexive (A) and voluntary (C) covert attention tasks are presented as a function of group (patients vs. controls) cue (valid vs. invalid) and stimulus onset asynchrony (SOA). Error bars represent the within-subject standard error (Loftus & Masson, 1994). **Boxplots for the cue effect size data** (i.e., invalid minus valid RTs) for the reflexive (B) and voluntary (D) covert attention tasks are presented as a function of group (patients vs. controls) and SOA. Error bars represent the interquartile range. For the boxplots, within each box X represents the mean and – represents the median. * indicates a statistically significant difference.

### Voluntary covert attention

For cued trials, a mixed-model ANOVA (group x cue x SOA (250, 350, 550)) revealed main effects of cue (F(1,32)=63.94, *p*<.001, η^2^_p_=.67) and SOA (F(2,64)=14.38, *p*<.001, η^2^_p_ =.31), with no other main effects or interactions (Figure 3C). The main effect of cue indicated that RTs were faster for valid (511ms) compared to invalid trials (554ms). The main effect of SOA indicated that the RTs for the 250ms SOA (545ms) and 350ms SOA (536ms) were significantly slower than RTs at the 550ms SOA (518ms, all *p*’s <.002, Holm-corrected). For comparison to the reflexive covert attention task, we also compared CES (invalid–valid RT) between the two groups at each SOA (Figure 3D). This analysis did not reveal any significant differences (all *p*’s >.20, uncorrected). Bayesian independent samples t-tests indicated anecdotal support for the null hypothesis for each SOA (all BF_10_ <0.42; see Supplementary Material).

### Non-spatial attention tasks

All accuracy and RT data for the non-spatial attention tasks for each patient and controls are presented in Supplementary Tables 3 & 4. For the non-spatial attention tasks, we collected data from 13 of 14 patients as one dropped out of the study before completing this session. One of the controls did not complete the non-spatial attention testing session resulting in a total of 23 controls for the AB and SART tasks. Of these remaining 23 controls, one had accuracy scores that were more than 2SD below the mean of controls for 2 different lags on the AB task and was removed as an outlier (see Methods). This resulted in 22 controls for the final analysis of the AB task and 23 controls for the final analysis of the SART.

### Attentional blink (AB). 1-Target trials

We first analyzed the percentage of errors for the 1-Target trials using a mixed-model ANOVA with lag (1, 2, 3, 4, 8 and 12) as the within-subject factor and group (patients vs. controls) as the between-subject factor. This analysis revealed no significant main effects or interactions (all *p*’s >.21). Thus, both groups performed equally well on 1-Target trials.

### 2-Target trials

To examine the AB effect, we analyzed 2-Target trials where both the response to target 1 (T1) and 2 (T2) were correct (Figure 4A). This mixed-model ANOVA with lag (1, 2, 3, 4, 8, 12) as a within-subject factor and group (patients vs. controls) as a between-subject factor revealed significant main effects of lag (F(5,165)=27.27, *p*<.001, η^2^_p_=.45), and group (F(1,33)=8.32, *p*=.007, η^2^_p_=.20; Figure 4B). Specifically, participants had higher accuracy in the last 3 lags (4, 8, 12) compared to the first 3 lags (1, 2, 3; all t’s >3.46, all *p*’s <.004, Holm-corrected). Patients also had lower accuracy overall for 2-Target trials (68%) compared to controls (78%). Although the group x lag interaction was not statistically significant (F(5,165)=1.67, *p*=.14, η^2^_p_=.05) a simple main effects analysis revealed that cerebellar patients performed more poorly than controls on 2-Target trials at lag-1 (patients=54% vs. controls=70%; t(33)=2.85, *p*=.04, Holm-corrected; d=1.0) and lag-3 (patients 58% vs. controls=72%; t(33)=2.98, *p*=.03, Holm-corrected; d=1.04). These results were further supported by Bayesian independent samples t-tests which indicated moderate support for the alternative hypothesis for both lag-1 (BF_10_=6.22) and lag-3 (BF_10_=8.14). All other BF_10_ <1.84 (see Supplementary Material).

**Figure 4.**
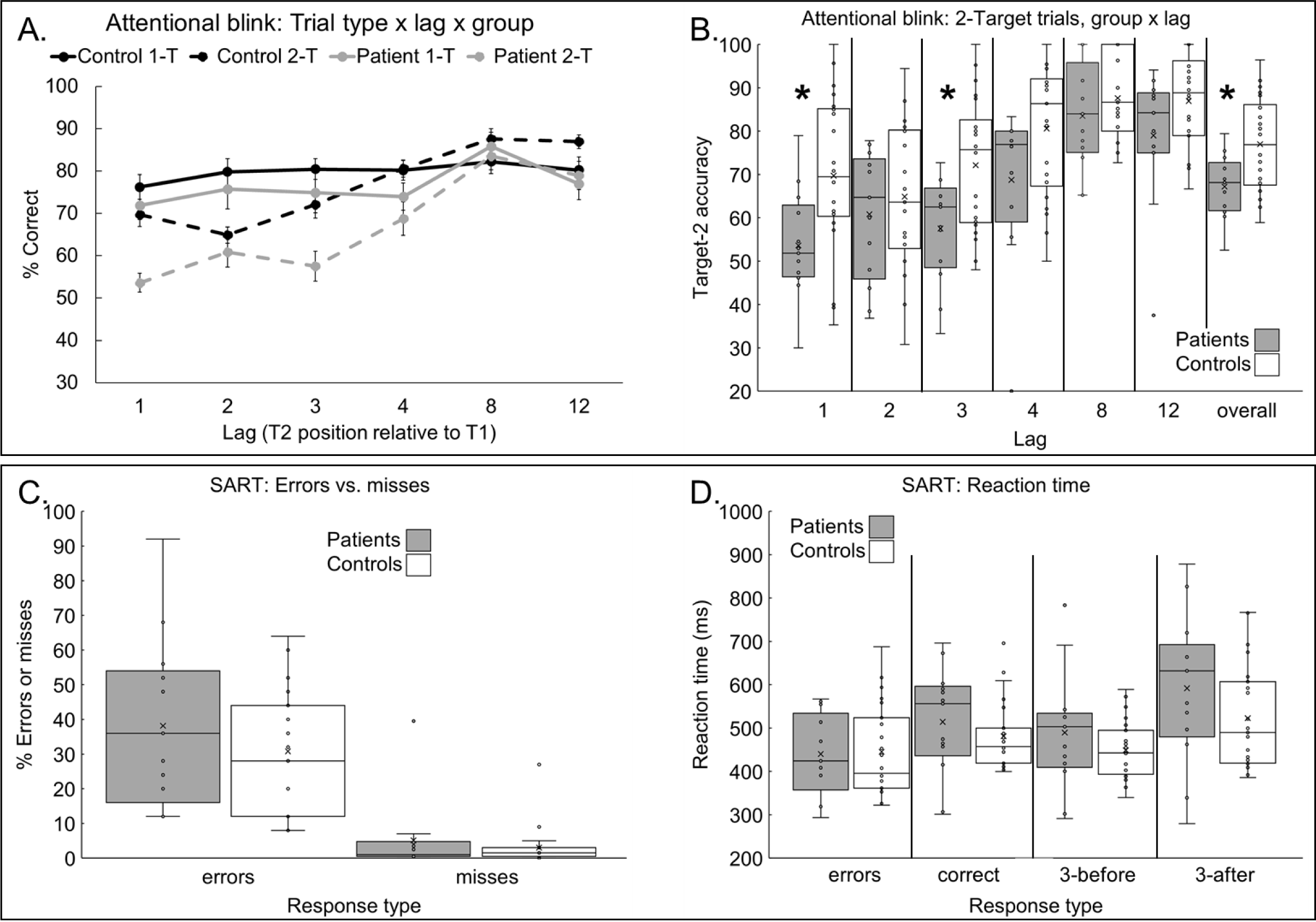
Accuracy data for the attention blink task (panel A) is presented as a function of group (patients vs. controls), lag (1, 2, 3, 4, 8, 12), and trial type (1-Target vs. 2-Target). Boxplot depicting the accuracy for 2-Target trials for the attentional blink task (panel B). Boxplots for the percentage of errors or misses (panel C) and reaction time (RT) data (panel D) for the sustained attention to response task (SART) are presented as a function of response type (hit, miss or error). Error bars for panel A represent the within-subject standard error (Loftus & Masson, 1994). Error bars for the remaining panels (B-D) represent the interquartile range. For the boxplots, within each box X represents the mean and – represents the median. * indicates a statistically significant difference.

### Lag-1 sparing

A number of previous studies investigating the AB have observed “lag-1 sparing” during 2-Target trials such that when T2 appears immediately after T1 the accuracy in reporting T2 is greater than when T2 appears at lag-2 or 3 (Dux & Marois, 2009; Raymond et al., 1992; Visser, Bischof, & Di Lollo, 1999). Lag-1 sparing is thought to occur if T2 appears immediately after T1 because both targets are processed as a single event or episode. According to Hommel & Akyurek (2005) and Visser et al. (Visser et al., 1999) lag-1 sparing is said to be present when accuracy at lag-1 is 5+% greater than the lag with the lowest level of performance. Alternatively, lag-1 sparing is said to be absent when performance at lag-1 is below the lowest level of performance at the next few lags. Controls were 5% more accurate at lag-1 (70%) compared to their lowest performance at lag-2 (65%). In contrast, patients showed the reverse effect with poorer performance at lag-1 (54%) compared to lags 2 (61%) and 3 (58%). Thus, according to the criteria outlined above, lag-1 sparing is relatively preserved in controls, but absent in cerebellar patients.

### Sustained attention to response task (SART)

For the SART we first compared the percentage of commission errors (i.e., presses for “3”) and misses (i.e., omission errors) across the two groups (Figure 4C). This analysis revealed no significant differences (commission errors; patients=38% vs. controls=31%; t(34)=1.058, *p*=.30; omission errors; patients=5% vs. controls=3%; t(34)=0.79, *p*=.44). A subsequent analysis using d’ also revealed no significant group differences in sensitivity (controls d’=2.73 vs. patients d’=2.36; t(34)=1.04, p=.31).

Next, we examined RTs by comparing the average RTs for errors of commission vs. correct responses between the two groups using a mixed-model ANOVA with trial type (errors vs. hits) as a within-subject factor and group (patients vs. controls) as a between-subject factor (Figure 4D). This analysis demonstrated a significant main effect of trial type with RTs for error trials (441ms) being faster than RTs for hits (496ms; F(1,34)=15.07, *p*<.001; η^2^_p_=.31). No other effects were significant.

We examined post error slowing using a similar ANOVA to compare the average RT for the three trials immediately preceding an error of commission compared to the average RTs for the three trials immediately following an error commission. This analysis also revealed a significant main effect of trial type such that participants were faster to respond in the 3 trials immediately preceding an error (462ms) compared to the 3 trials following an error (550ms; F(1,34)=30.29, *p*<.001; η^2^_p_ =.47). No other effects were significant.

To examine RT variability, we conducted a trial type x group ANOVA on the SD of RTs for errors of commission, as well as RTs for correct trials. This analysis revealed no main effects or interactions, indicating that RT variability was not different between cerebellar patients and controls (all *p*’s >.21).

Finally, Bayesian independent samples t-tests revealed anecdotal support for the null hypothesis for errors of commission (BF_10_=0.51), misses (BF_10_=0.42), d-prime (BF_10_=0.50), as well as RTs for errors (BF_10_=0.33) and correct trials (BF_10_=0.48; Supplementary Material).

### Correlation analysis

In addition to analyzing data from the attention tasks, we also wanted to examine whether performance on any of the attention tasks was correlated with any relevant clinical measures such as time post stroke, lesion volume, or motor impairment (i.e., total ICARS score). No significant correlations emerged between overall CES for the two covert attention tasks or performance on the AB (2-Target accuracy) or SART (errors of commission) tasks with either lesion volume, time post stroke, or the overall score on the ICARS (all *p*’s >.073, uncorrected).

### Lesion data

Lesion maps for each patient are available online (Supplementary Figure 1). The coordinates of the area of maximum lesion overlap in our group (in red) is the inferior semilunar lobule (i.e., Crus II; MNI: x= −22, y= −74, z= −48; MNI: x= −17, y=-76, z= −45). This region was damaged in 10 of the 14 patients (i.e., 70% overlap; Figure 5).

**Figure 5.**
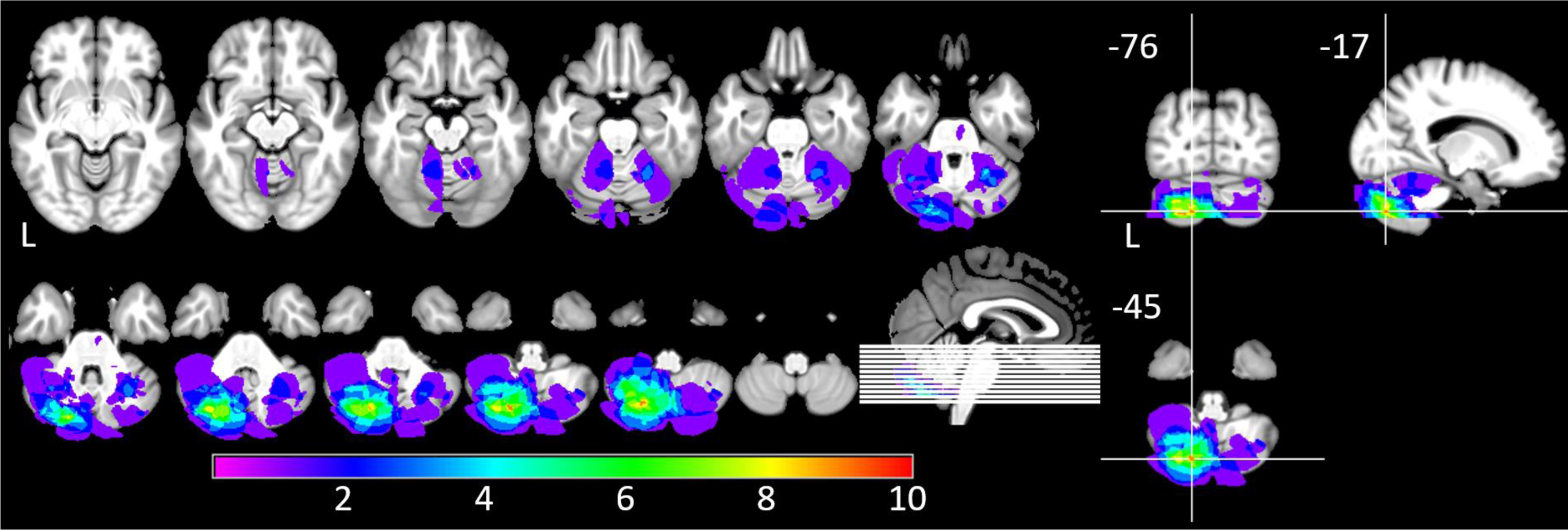
Lesion overlap for the cerebellar patient group (n=14). The region of maximum lesion overlap is the inferior semilunar lobule (i.e., Crus II; MNI: x= −17, y= −76, z= −45 and x= −22 y=-74, z= −48) in the posterior lobe of the left cerebellum. The legend indicates the number of patients in the sample that have damage to a given region. The regions with the largest degree of overlap were damaged in at least 10/14 patients (i.e., 70% overlap; depicted in red).

In subsequent analyses we used voxel-based lesion symptom mapping (VLSM; see Methods) to examine the relationship between cerebellar damage and increased CES at the 600ms SOA in the reflexive covert attention task, as well as poorer accuracy for 2-Target trials on the AB. For the reflexive covert attention task this analysis revealed a significant relationship between damage in the left inferior semilunar lobule (i.e., Crus II; MNI: X=-24, Y=-71, Z=-44) and increased CES in patients (Brunner-Munzel Z-test, *p*<.05, uncorrected; Supplementary Figure 2).

Similar to the analysis with the covert attention data, the VLSM analysis of the AB data indicated that the damaged region that was most associated with impaired accuracy on 2-Target trials, was, again, the left inferior semilunar lobule (i.e., Crus II; X=-24, Y=-71, Z=-44; Brunner-Munzel Z-test, p<.05, uncorrected; Supplementary Figure 3).

It is important to note that, although both VLSM analyses support our lesion overlay data, these results do not survive correction for multiple comparisons due to the small number of patients in our sample. As such, these results are presented in the Supplementary Material, but should be interpreted with caution.

Finally, to check for evidence of damage to the dentate nucleus we localized the dentate using the probabilistic 3D atlas developed by Dimitrova and colleagues (2006). Based on their maximal MNI coordinates for the left (X= −15, Y= −57, Z= −36) and right (X= 19, Y= −55, Z= −36) dentate, only two patients had damage to the dentate nucleus (patient 909, left dentate; patient 523, probable right dentate). Note that the significant difference in cueing effect at the 600ms SOA in the reflexive covert attention task and the increased AB effect are still apparent in our cerebellar patients even when the two patients with dentate damage are excluded from the analyses (see Supplementary Material).

## Discussion

Cerebellar damage impairs the ability to orient attention in both time and space. Specifically, for reflexive orienting of spatial attention we demonstrated clear evidence that cerebellar damage results in a significantly larger cueing effect at the longest (i.e., 600ms) SOA which may reflect a slowed onset of IOR (Figure 3A&B). Specifically, the cueing effect for cerebellar patients was still quite large (54ms) compared to a non-significant cueing effect for controls (−5ms; Figure 3B). The increased cueing effect in cerebellar patients at the longest (600ms) SOA provides a conceptual replication and extension of our earlier work (Striemer, Cantelmi, et al., 2015) in which patients with cerebellar injury showed a trend towards a slowed onset of IOR at a shorter (300ms) SOA. However, in contrast to our previous study (Striemer, Cantelmi, et al., 2015), we did not see any evidence of a slower orienting response at the earliest (50ms) SOA. Although it is unclear why this effect at the earliest SOA was not observed here, the slowed onset of IOR appears to be a consistent impairment observed following damage to the lateral cerebellum as this effect has now been observed in two separate studies with different patient groups. Previous studies suggest that IOR may be related to an inhibition of response to a previously attended location (for reviews see Chica, Martin-Arevalo, Botta, & Lupianez, 2014; Klein, 2000). Thus, damage to the posterior-lateral cerebellum may impair the ability to inhibit responding at previously attended locations. Confirmation of this hypothesis will require future investigation.

It is important to note that, although patients showed still showed a significant cueing effect at the 600ms SOA the cueing effect for controls was not different from zero. Typically, a significant IOR effect (i.e., a negative cueing effect) is observed at SOAs >250ms in covert attention tasks that use non-predictive peripheral cues (Chica et al., 2014; Posner et al., 1985). One of the reasons it may have been more difficult for us to observe IOR in our control group is that we used a reflexive covert attention task that included cue-target overlap. Typically, cue-target overlap can slow the onset of IOR in healthy adults (Collie, Maruff, Yucel, Danckert, & Currie, 2000). We chose to use cue target overlap to ensure that we would observe significant cueing effects at early SOAs (Collie et al., 2000) and to make our results directly comparable to our previous work with cerebellar patients (Striemer, Cantelmi, et al., 2015). We suggest that the increased cueing effect we observed at the 600ms SOA in cerebellar patients is evidence of a slowed onset of IOR because this is the natural tendency at longer SOAs in reflexive covert attention tasks (Chica et al., 2014). Future research could either eliminate the cue-target overlap used here, or test at longer SOAs to ensure a direct comparison with a healthy control group exhibiting a significant IOR effect.

In the current study voluntary covert attention appeared relatively unaffected by cerebellar damage. This result is consistent with earlier work suggesting that the cerebellum may play less of a role in voluntary attention (Oliver Baumann & Mattingley, 2014; Striemer, Chouinard, et al., 2015). In fact, a previous fMRI study from our lab observed greater BOLD activation in the left cerebellum for reflexive (i.e., peripheral cue) compared to voluntary (i.e., central arrow cue) covert attention tasks in healthy adults (Striemer, Chouinard, et al., 2015). This greater impairment in reflexive attention may be due to the fact that peripheral abrupt onset cues (like the ones used in the reflexive attention task in the current study) are effective in attracting both shifts of attention (Jonides & Yantis, 1988), as well as eye movements (Mulckhuyse, van Zoest, & Theeuwes, 2008; Schreij, Owens, & Theeuwes, 2008). In contrast, central arrow cues require more time to be interpreted before a saccade plan can be generated (Jonides, 1981; Muller & Rabbitt, 1989). Therefore, it is likely that abrupt onset peripheral cues result in a stronger engagement of oculomotor structures in both the cerebral cortex and cerebellum thereby making these trials more likely to be affected by cerebellar injury. Although we did not observe any significant effects of cerebellar lesions on voluntary covert attention, we cannot rule out the possibility that a significant effect might be detected in future studies using a larger patient sample. This is supported by the fact that Bayesian t-tests indicated only anecdotal support for the null hypothesis for the voluntary covert attention task (see Supplementary Material).

It is also important to acknowledge that previous work by Ristic and Kingstone and others (Hommel, Pratt, Colzato, & Godijn, 2001; Ristic, Friesen, & Kingstone, 2002; Ristic & Kingstone, 2006) has argued that central arrow cues may also trigger reflexive shifts of attention. Thus, one could argue that our voluntary covert attention task was also indexing reflexive attention. However, in the current study we used *predictive* arrow cues (70% valid) which participants were told about ahead of time. Thus, even if one suggests that arrow cues may lead to an initial reflexive shift of attention, the predictive nature of the cues used in the current study would lead to an additional voluntary shift of attention (Ristic & Kingstone, 2006), especially at the longer SOAs used in the current experiment (250+ ms).

Cerebellar damage also resulted in impaired performance on the AB, a task that is commonly used to measure “temporal attention” – or one’s capacity to allocate attention to specific moments in time (Dux & Marois, 2009; Schweizer et al., 2007). That is, cerebellar damage resulted in poorer overall performance on the 2-Target trials for patients vs. controls (Figure 4 A&B). In addition, according to the criteria outlined by Hommel and Akyurek (2005) and Visser and colleagues (1999), the well-known lag-1 sparing effect was absent for cerebellar patients but was present in controls. Previous research suggests that lag-1 sparing is associated with the binding of the two targets into a single event when they are presented in close temporal proximity without an intervening distractor (Dux & Marois, 2009; Hommel & Akyurek, 2005; Raymond et al., 1992). Thus, the absence of lag-1 sparing and a larger overall AB effect in cerebellar patients implies a slowed rate of processing each item in the RSVP stream. This is consistent with the notion that the cerebellum plays an important role in the detection and processing of sequences (O. Baumann et al., 2015; Leggio & Molinari, 2015), as well as anticipating the timing of sensory events in order to reduce performance variability (Ghajar & Ivry, 2009). Damage to such a mechanism would make it difficult to process a rapid series of events over a short time scale as is required during a task like the AB.

Although cerebellar damage resulted in problems with the processing of a rapid series of visual events over time as quantified by the AB, there was no problem with sustained attention per se as cerebellar patients performed similarly to controls on the SART. Specifically, patient’s error rates, sensitivity, RTs, RT variability and post-error slowing on the SART were statistically indistinguishable from controls, suggesting that the cerebellar regions damaged in our patients are not involved in updating performance following attentional errors. If one assumes a vigilance task like the SART requires some degree of performance monitoring (an executive function, Jurado & Rosselli, 2007) then the absence of any impairment on the SART following cerebellar injury is somewhat surprising given that previous research has argued that the cerebellum plays an important role in performance monitoring (Peterburs & Desmond, 2016) and executive functions more generally (Bellebaum & Daum, 2007). Consistent with the present findings, two previous patient studies observed that cerebellar lesions did not impair performance on a Go/NoGo attention task that is similar in nature to the SART (Gottwald, Mihajlovic, Wilde, & Mehdorn, 2003; Gottwald, Wilde, Mihajlovic, & Mehdorn, 2004). However, neuroimaging studies tell a different story. Specifically, neuroimaging studies suggest that Crus I in the left cerebellum is consistently activated during tasks that measure executive functions (see Stoodley & Schmahmann, 2009b for a meta-analysis). It is worth noting that the region of maximum lesion overlap in the current study was Crus II (Figure 5). Thus, the fact that we did not observe deficits in SART task performance might be related to the fact that, on average, Crus I was relatively spared in our patients. Again, it is important to note that, although we did not observe any effects of cerebellar lesions on the SART this should be further verified in a larger patient sample as Bayesian t-tests provided only anecdotal support for the null hypothesis.

As mentioned previously, our lesion data revealed that the region most consistently damaged in our patients was Crus II (inferior semilunar lobule) of the left posterior cerebellum (Figure 5). This observation was further supported by voxel-based lesion symptom mapping (VLSM) analyses which indicated that damage to Crus II was related to impaired performance on the reflexive covert attention task and the AB (see Supplementary Figures 2 & 3). However, these VLSM data should be interpreted with caution due to our smaller patient sample.

Finally, we also examined each patient’s scan for any evidence of damage to the cerebellar dentate nucleus with only two patients showing evidence of damage to this region (in opposing hemispheres). This result is important given that damage to the dentate nucleus would effectively disconnect the entire lateral cerebellar hemisphere from the cerebral cortex. Thus, the attention deficits observed here are due to damage to Crus II of the left cerebellum, and not the disruption of an entire cerebellar hemisphere. This was further reinforced by follow-up analyses (see Supplementary Material) where the reduced IOR effect and the increased AB effect were still present in our cerebellar patients even when the data from the two patients with dentate damage were removed. It is important to note that resting state functional connectivity data indicate that the same cerebellar regions damaged in our patients are functionally connected with the dorsal and ventral attention networks (Brissenden, Levin, Osher, Halko, & Somers, 2016; Buckner et al., 2011; Guell, Schmahmann, Gabrieli, & Ghosh, 2018; Wang et al., 2013) and are active during divided attention tasks (King, Hernandez-Castillo, Poldrack, Ivry, & Diedrichsen, 2018).

It is noteworthy that the lesion overlap and VLSM data indicated that the attentional deficits we observed following cerebellar lesions were most apparent after damage to the *left* cerebellum. Again, this is consistent with our previous fMRI study (Striemer, Chouinard, et al., 2015) demonstrating significant BOLD activity in the left cerebellum during covert attention tasks. In addition, an earlier fMRI study by Allen and colleagues (Allen, Buxton, Wong, & Courchesne, 1997) also observed significant BOLD activity in the left cerebellum when participants completed a non-spatial attention task after motor responses were controlled for. Furthermore, resting state functional connectivity data from Bucker and colleagues has revealed that that regions of the left cerebellum such as lobule VI, VIII and Crus II constitute the most strongly lateralized network in the left cerebellum. Importantly, this left lateralized cerebellar network is functionally connected to the ventral attention network in the right hemisphere (Buckner et al., 2011; Wang et al., 2013) which is known to play a critical role in covert attention, as well as temporal attention, and target detection (Corbetta, Kincade, Ollinger, McAvoy, & Shulman, 2000; Corbetta & Shulman, 2002; Husain & Nachev, 2007; Husain et al., 1997).

One limitation of the current study is that, of the patients that we were able to recruit (n=14), 8 had lesions the left cerebellum, 4 had bilateral lesions, and only 2 had isolated right cerebellar lesions. This makes it difficult to directly compare the effects of left vs. right cerebellar lesions on attention. This could be addressed in future studies by employing the same attention tasks in patients with right cerebellar lesions, or by using non-invasive brain stimulation techniques (e.g., TMS or tDCS) to examine the effects of stimulation of the left or right cerebellum on attention.

It important to emphasize that the attentional deficits observed in our cerebellar patients are not simply due to motor impairments. Specifically, there were no significant correlations between the total score on the ICARS and performance measures on any of the attention tasks that we used. In addition, although patient RTs were somewhat slower overall than controls for the reflexive covert attention task (*p*=.058), there were no significant differences in RT between the groups for the voluntary covert attention task, or any of the RT measures in the SART. Even if there was a general slowing of responses in our cerebellar patients (of which there is limited evidence) it would still not explain the main findings in the current study. Specifically, for the reflexive covert attention task, we see a significantly larger cueing effect at the 600ms SOA in patients compared to controls. It is important to point out that cueing effects are calculated by subtracting the RT for valid trials from the RT for invalid trials. This difference in RT indexes the effectiveness of the cue, while controlling for overall response speed between the two conditions. Thus, a general slowing of responses cannot explain differences in cueing effects because a general slowing in RT would affect valid and invalid trials equally. Alternatively, for the AB task, we measured the accuracy with which participants report the presence of targets. The decreased accuracy for the 2-Target task for patients compared to controls cannot be explained by a general slowing of responses because this measure is not dependent on RT.

### What role does the cerebellum play in attention?

Our findings support the notion that cerebellar damage disrupts the allocation of attention across spatial and temporal domains resulting in a form of “attentional dysmetria.” One prominent theory suggests that the cerebellum coordinates motor output by comparing it to predicted sensory consequences (Ghajar & Ivry, 2009; Sokolov, Miall, & Ivry, 2017). The cerebellum may then “generate time-based expectancies of sensory information” in order to more efficiently synchronize predicted with actual sensory input in order to help reduce performance variability (Ghajar & Ivry, 2009). This prediction is similar to suggestions that lesions to the cerebellum produce “dysmetria of thought” through disrupting the timing and coordination of cognitive processing (Schmahmann et al., 2019; Schmahmann & Sherman, 1998). Our data directly support these theories by demonstrating that lesions to the same cerebellar region (left Crus II) leads to an impairment on the AB (see also Schweizer et al., 2007), as well as the slowed onset of IOR. The fact that lesions to the same region of the cerebellum can disrupt both “spatial” and “temporal” attention is perhaps not surprising if one considers attention from a limited capacity resource perspective where both spatial and temporal attention are thought to rely on similar underlying cognitive and neural mechanisms (e.g., Dell’Acqua, Sessa, Jolicoeur, & Robitaille, 2006; Husain & Nachev, 2007; Malhotra, Coulthard, & Husain, 2009).

In summary, we have demonstrated, for the first time, that damage to Crus II in the left posterior-lateral cerebellum results in a significantly larger cueing effect at longer SOAs which may indicate a slowed onset of IOR, and reduces the capacity to detect successive targets during a rapid serial visual presentation (i.e., an increased AB effect). Importantly, anatomical studies in non-human primates, as well as connectivity and task-based functional neuroimaging studies in humans, have demonstrated that this same cerebellar region is linked with cortical networks that are known to be involved in attention. These data therefore provide direct support for the notion that the cerebellum plays a critical role in both spatial and temporal attention.

## Acknowledgements

The authors would like to thank Nadia Botha, Suekiana Choucair and Hanbin Go for their assistance with data collection and analysis, and Drs. Jörn Diedrichsen and Carlos Hernandez-Castillo (Brain and Mind Institute, University of Western Ontario) for their assistance with the cerebellar lesion analysis. The authors would also like to thank Nadine Quehl and Karly Neath-Tavares for their assistance with patient recruitment. This research was supported by Natural Sciences and Engineering Research Council of Canada (NSERC) Discovery Grants to C.S. (2014-04542) & J.D. (50503-10762), a Canadian Foundation for Innovation (CFI) grant to J.D. (52046-10013), and a Strategic Research Grant (01229) from MacEwan University to C.S.

## Supplementary material

### 1. Bayesian independent samples t-tests for the covert attention, attentional blink, and sustained attention to response tasks

To examine the strength of the evidence in favor of the null or the alternative hypothesis for effects of interest for each task we computed Bayesian independent samples t-test using JASP (JASP-Team, 2020). We present inverse Bayes factors BF_10_ where larger values (i.e., >1.0) mean more support for the alternative hypothesis (i.e., H_1_) and smaller values (i.e., <1.0) mean more support for the null hypothesis (i.e., H_0_; Jarosz & Wiley, 2014; Masson, 2011; Wagenmakers, 2007).

#### Reflexive covert attention

Between group comparison of cue-effect size at each SOA

**Reflexive covert attention:** Between group comparison of cue-effect size at each SOA **Bayesian Independent Samples T-Test**
|  | **$BF_{10}$** | **evidence** |
| --- | --- | --- |
| CES_50 | 0.497 | $H_0$ = anecdotal |
| CES_100 | 0.553 | $H_0$ = anecdotal |
| CES_300 | 0.797 | $H_0$ = anecdotal |
| CES_600 | 22.49 | $H_1$ = strong |

#### Voluntary covert attention

Between group comparison of cue-effect size at each SOA

**Voluntary covert attention:** Between group comparison of cue-effect size at each SOA **Bayesian Independent Samples T-Test**
|  | **$BF_{10}$** | **evidence** |
| --- | --- | --- |
| CES_250 | 0.417 | $H_0$ = anecdotal |
| CES_350 | 0.544 | $H_0$ = anecdotal |
| CES_550 | 0.655 | $H_0$ = anecdotal |

#### Attentional Blink (AB)

Between groups comparison of performance on T2 trials at each lag.

**Attentional Blink (AB):** Between groups comparison of performance on T2 trials at each lag. **Bayesian Independent Samples T-Test**
|  | **$BF_{10}$** | **evidence** |
| --- | --- | --- |
| T2_lag_1 | 6.221 | $H_1$ = moderate |
| T2_lag_2 | 0.409 | $H_0$ = anecdotal |
| T2_lag_3 | 8.143 | $H_1$ = moderate |
| T2_lag_4 | 1.835 | $H_1$ = anecdotal |
| T2_lag_8 | 0.537 | $H_0$ = anecdotal |
| T2_lag_12 | 1.175 | $H_1$ = anecdotal |

#### Sustained attention to response task (SART)

Between group comparisons of errors of commission (errors_3), errors of omission (perc_miss), d-prime, RT for errors of commission (3press_RT), RT for correct responses (correct_rt), RTs for the 3 trials preceding (Prior RT) and 3 following (Post RT) an error, and the SD of the RTs for errors of commission (3press STDEV) and correct responses (correct STDEV).

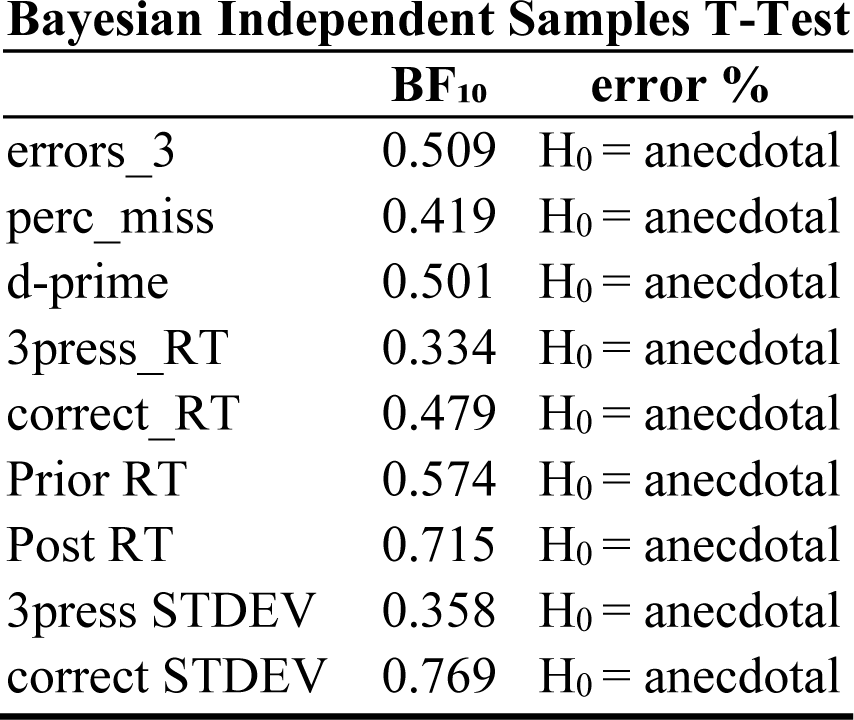

**Bayesian Independent Samples T-Test**
|  | **BF<sub>10</sub>** | **error %** |
| --- | --- | --- |
| errors_3 | 0.509 | H<sub>0</sub> = anecdotal |
| perc_miss | 0.419 | H<sub>0</sub> = anecdotal |
| d-prime | 0.501 | H<sub>0</sub> = anecdotal |
| 3press_RT | 0.334 | H<sub>0</sub> = anecdotal |
| correct_RT | 0.479 | H<sub>0</sub> = anecdotal |
| Prior RT | 0.574 | H<sub>0</sub> = anecdotal |
| Post RT | 0.715 | H<sub>0</sub> = anecdotal |
| 3press STDEV | 0.358 | H<sub>0</sub> = anecdotal |
| correct STDEV | 0.769 | H<sub>0</sub> = anecdotal |

### 2. Voxel-based lesion symptom mapping analyses

As a supplement to our lesion overlay data in the main manuscript we also carried out voxel-based lesion symptom mapping (VLSM) with the NPM toolbox in MRIcron (Rorden, Karnath, & Bonilha, 2007). This analysis allowed us to examine whether damage to specific regions of the cerebellum were linked with impaired performance on the reflexive covert attention task (i.e., a larger cueing effect at the 600ms SOA). The results of this analysis revealed that the damaged region that had the highest association with impaired performance on the reflexive covert attention task was the inferior semilunar lobule (i.e., Crus II; X=-24, Y=-71, Z=-44) of the left lateral cerebellum, (Brunner-Munzel Z-test, *p*<.05, uncorrected; Supplementary Figure 2).

In addition, we also used VLSM to examine the damaged regions of the cerebellum that were associated with poorer performance on the AB task (i.e., decreases in T2 accuracy). Similar to the analysis with the covert attention data, this analysis indicated that the damaged region that was most associated with impaired T2 accuracy was the inferior semilunar lobule (i.e., Crus II; X=-24, Y=-71, Z=-44) of the left lateral cerebellum (Brunner-Munzel Z-test, *p*<.05, uncorrected; Supplementary Figure 3).

It is important to note that, although both VLSM analyses support our initial conclusions, these results are presented with uncorrected *p*-values as the analyses do not survive correction for multiple comparisons (FDR or Bonferroni) due to the small number of patients. As such, these results should be interpreted with caution.

### 3. Attentional deficits are not the result of damage to the dentate nucleus

To ensure that the attentional deficits we observed in the reflexive covert attention task and the AB task were not related to damage to the dentate nucleus we re-ran the same analyses again not including the data from the two patients with probable dentate damage (patients 523 and 909; see Table 2 in the main manuscript).

Re-analysis of the data from the reflexive covert attention task revealed identical results. Specifically, there was a significant cue x SOA x group interaction (F(3,93)=3.47, *p*=.019; η^2^_p_=.10) such that CES for the cerebellar patients was larger at the 600ms SOA (patients=54ms vs. controls=-2ms; (t(31)=3.26, *p*=.012, Holm corrected). However, there were no significant differences at any of the other SOAs (all *p*’s >.11, uncorrected).

In addition, we also re-analyzed the AB data removing the two patients with dentate damage. This analysis produced identical results with a main effect of group (F(1,31)=5.84, *p*=.021, η^2^_p_=.16) indicating significantly poorer performance on 2-Target trials for patients (70%) vs. controls (78%).

---

**Supplementary Table 1:** Mean reaction time (RT) data for the reflexive covert attention task for patients (n=11) and controls (n=23) as a function of cue (valid vs. invalid) and stimulus onset asynchrony (50, 100, 300, 600ms).

| **Patient (lesion):** | **Valid 50** | **Valid 100** | **Valid 300** | **Valid 600** | **Invalid 50** | **Invalid 100** | **Invalid 300** | **Invalid 600** |
| --- | --- | --- | --- | --- | --- | --- | --- | --- |
| 29 (L) | 965 | 930 | 914 | 902 | 1009 | 980 | 1094 | 1104 |
| 61 (L) | 509 | 513 | 529 | 549 | 582 | 553 | 576 | 582 |
| 182 (L) | 613 | 529 | 661 | 552 | 619 | 649 | 573 | 660 |
| 309 (L) | 576 | 541 | 482 | 490 | 627 | 584 | 481 | 465 |
| 378 (R) | 454 | 459 | 490 | 490 | 497 | 507 | 470 | 480 |
| 522 (L) | 563 | 494 | 501 | 485 | 613 | 589 | 567 | 605 |
| 523 (R) | 606 | 566 | 528 | 488 | 621 | 597 | 595 | 547 |
| 564 (L) | 610 | 593 | 592 | 600 | 677 | 699 | 652 | 699 |
| 678 (L) | 459 | 454 | 454 | 463 | 474 | 481 | 444 | 436 |
| 867 (B) | 593 | 579 | 583 | 613 | 659 | 690 | 658 | 628 |
| 953 (L) | 719 | 676 | 674 | 729 | 730 | 785 | 816 | 750 |
| **Patient mean** | **606** | **576** | **582** | **578** | **646** | **647** | **630** | **632** |
| **Patient SD** | **141** | **134** | **131** | **133** | **141** | **141** | **186** | **185** |
| **Control mean** | **523** | **514** | **521** | **536** | **572** | **570** | **542** | **533** |
| **Control SD** | **65** | **72** | **75** | **70** | **64** | **76** | **77** | **82** |

**Supplementary Table 2:** Mean reaction time (RT) data for the voluntary covert attention task for patients (n=11) and controls (n=23) as a function of cue (valid vs. invalid) and stimulus onset asynchrony (250, 350, 550ms).

| **Patient (lesion):** | **Valid 250** | **Valid 350** | **Valid 550** | **Invalid 250** | **Invalid 350** | **Invalid 550** |
| --- | --- | --- | --- | --- | --- | --- |
| 29 (L) | 731 | 787 | 782 | 931 | 840 | 821 |
| 61 (L) | 442 | 440 | 432 | 479 | 481 | 493 |
| 182 (L) | 549 | 543 | 591 | 586 | 615 | 584 |
| 309 (L) | 397 | 349 | 319 | 428 | 382 | 395 |
| 378 (R) | 445 | 440 | 453 | 469 | 469 | 462 |
| 522 (L) | 532 | 555 | 464 | 556 | 620 | 579 |
| 523 (R) | 604 | 526 | 569 | 592 | 590 | 556 |
| 564 (L) | 587 | 579 | 563 | 724 | 640 | 745 |
| 678 (L) | 408 | 361 | 321 | 411 | 412 | 313 |
| 867 (B) | 670 | 627 | 653 | 683 | 685 | 659 |
| 953 (L) | 656 | 627 | 561 | 640 | 779 | 633 |
| **Patient mean** | **547** | **530** | **519** | **591** | **592** | **567** |
| **Patient SD** | **114** | **129** | **139** | **152** | **146** | **148** |
| **Control mean** | **515** | **497** | **488** | **548** | **540** | **515** |
| **Control SD** | **77** | **77** | **83** | **87** | **99** | **86** |

**Supplementary Table 3:** Mean percent accuracy on the attentional blink task for patients (n=13) and controls (n=22) as a function of trial type (1-Target vs. 2-Target) and Lag (1-4, 8 or 12).

| **Patient (lesion):** | **1T-L1** | **1T-L2** | **1T-L3** | **1T-L4** | **1T-L8** | **1T-L12** | **2T-L1** | **2T-L2** | **2T-L3** | **2T-L4** | **2T-L8** | **2T-L12** |
| --- | --- | --- | --- | --- | --- | --- | --- | --- | --- | --- | --- | --- |
| 29 (L) | 85 | 100 | 50 | 92 | 69 | 60 | 47 | 37 | 50 | 63 | 88 | 85 |
| 61 (L) | 43 | 67 | 75 | 62 | 40 | 83 | 30 | 72 | 63 | 20 | 100 | 76 |
| 117 (B) | 67 | 50 | 100 | 70 | 87 | 85 | 65 | 71 | 69 | 77 | 84 | 63 |
| 182 (L) | 92 | 100 | 100 | 81 | 75 | 67 | 55 | 75 | 65 | 77 | 92 | 89 |
| 309 (L) | 67 | 43 | 67 | 85 | 100 | 60 | 47 | 38 | 73 | 70 | 100 | 80 |
| 378 (R) | 71 | 90 | 20 | 88 | 100 | 79 | 68 | 60 | 63 | 80 | 65 | 94 |
| 523 (R) | 44 | 63 | 50 | 56 | 78 | 33 | 44 | 44 | 69 | 56 | 65 | 38 |
| 564 (L) | 82 | 75 | 64 | 57 | 89 | 89 | 46 | 48 | 39 | 78 | 76 | 84 |
| 670 (B) | 86 | 73 | 86 | 77 | 100 | 83 | 61 | 65 | 63 | 83 | 84 | 88 |
| 678 (L) | 50 | 100 | 100 | 71 | 92 | 88 | 79 | 72 | 57 | 76 | 100 | 92 |
| 867 (B) | 91 | 88 | 71 | 89 | 100 | 91 | 53 | 54 | 47 | 54 | 80 | 88 |
| 909 (B) | 70 | 75 | 91 | 65 | 100 | 100 | 50 | 77 | 58 | 80 | 74 | 75 |
| 953 (L) | 88 | 62 | 100 | 71 | 86 | 83 | 52 | 78 | 33 | 80 | 78 | 75 |
| **Patient mean** | **72** | **76** | **75** | **74** | **86** | **77** | **54** | **61** | **58** | **69** | **84** | **79** |
| **Patient SD** | **17** | **19** | **25** | **12** | **17** | **18** | **12** | **15** | **12** | **18** | **12** | **15** |
| **Control mean** | **76** | **80** | **80** | **80** | **82** | **80** | **70** | **65** | **72** | **81** | **88** | **87** |
| **Control SD** | **16** | **17** | **15** | **13** | **17** | **16** | **18** | **16** | **15** | **15** | **9** | **11** |

**Supplementary Table 4:** Mean percentage of errors and misses and mean reaction times (RT) for errors, hits, as well as the three trials preceding and following an error for patients (n=13) and controls (n=23) for the sustained attention to response task (SART).

| **Patient (lesion):** | **% errors** | **% misses** | **error RT** | **Hits RT** | **Errors:**<br>**3 Prior RT** | **Errors:**<br>**3 Post RT** |
| --- | --- | --- | --- | --- | --- | --- |
| 29 (L) | 48 | 39.5 | 567 | 696 | 691 | 878 |
| 61 (L) | 36 | 3.5 | 561 | 475 | 525 | 497 |
| 117 (B) | 56 | 4.5 | 555 | 673 | 784 | 827 |
| 182 (L) | 36 | 0.5 | 319 | 415 | 418 | 462 |
| 309 (L) | 92 | 5 | 294 | 301 | 291 | 280 |
| 378 (R) | 68 | 0.5 | 324 | 307 | 302 | 339 |
| 523 (R) | 12 | 1 | 469 | 603 | 543 | 665 |
| 564 (L) | 24 | 7 | 473 | 556 | 505 | 638 |
| 670 (B) | 28 | 2.5 | 391 | 590 | 435 | 632 |
| 678 (L) | 52 | 1 | 408 | 457 | 458 | 558 |
| 867 (B) | 12 | 1 | 424 | 564 | 509 | 720 |
| 909 (B) | 20 | 0.5 | 514 | 582 | 503 | 664 |
| 953 (L) | 12 | 0 | 424 | 465 | 400 | 536 |
| **Patient mean** | **38** | **5** | **440** | **514** | **490** | **592** |
| **Patient SD** | **24** | **11** | **94** | **125** | **136** | **173** |
| **Control mean** | **31** | **3** | **445** | **481** | **449** | **523** |
| **Control SD** | **17** | **6** | **103** | **78** | **67** | **119** |

**Supplementary Figure 1 Part 1.**
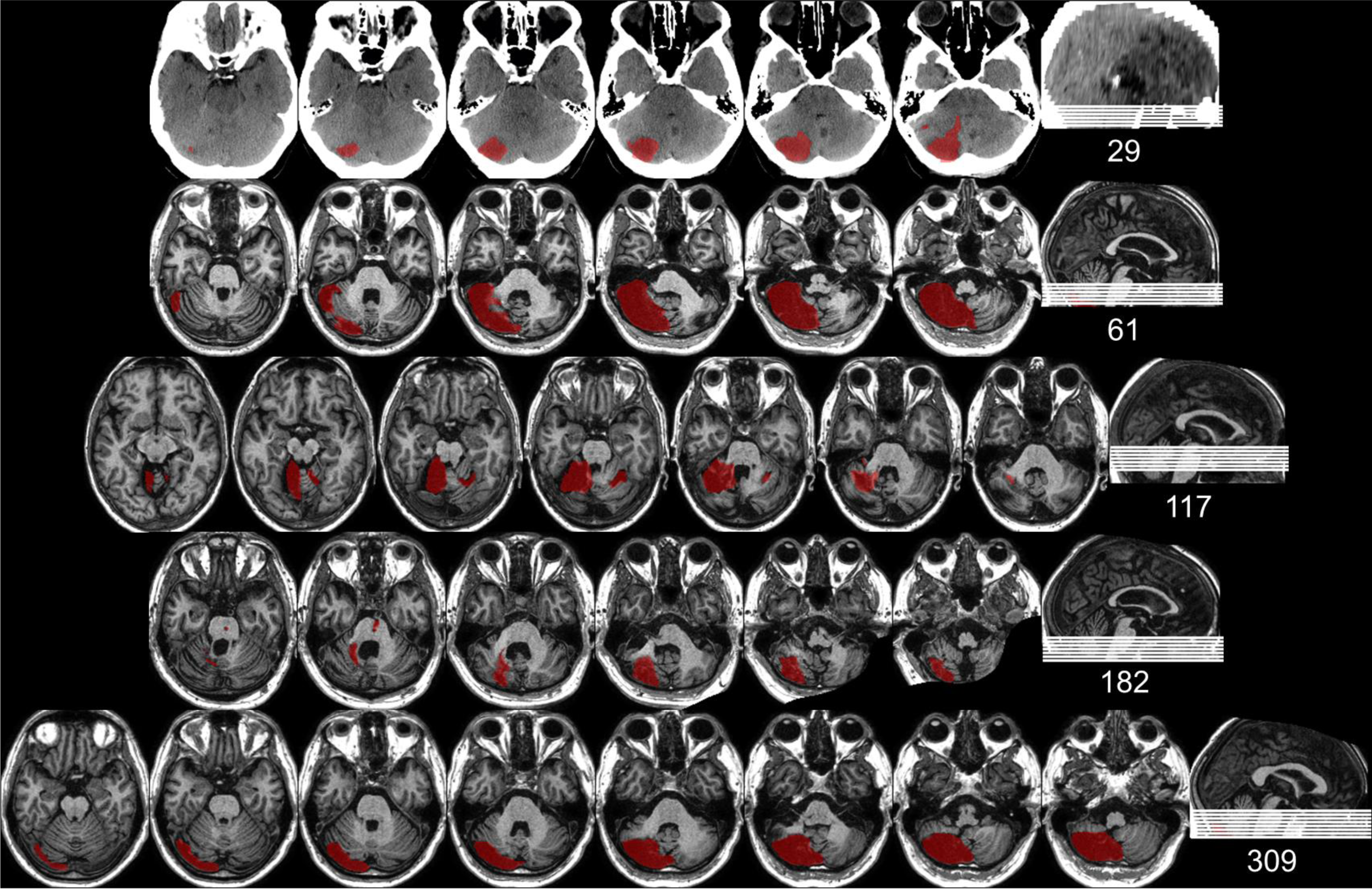
Individual lesion maps for each of the patients (patient number located to the right). See Table 2 in the main manuscript for clinical details for each patient. Images are presented in neurological convention (i.e., left is on the left).

**Supplementary Figure 1 Part 2.**
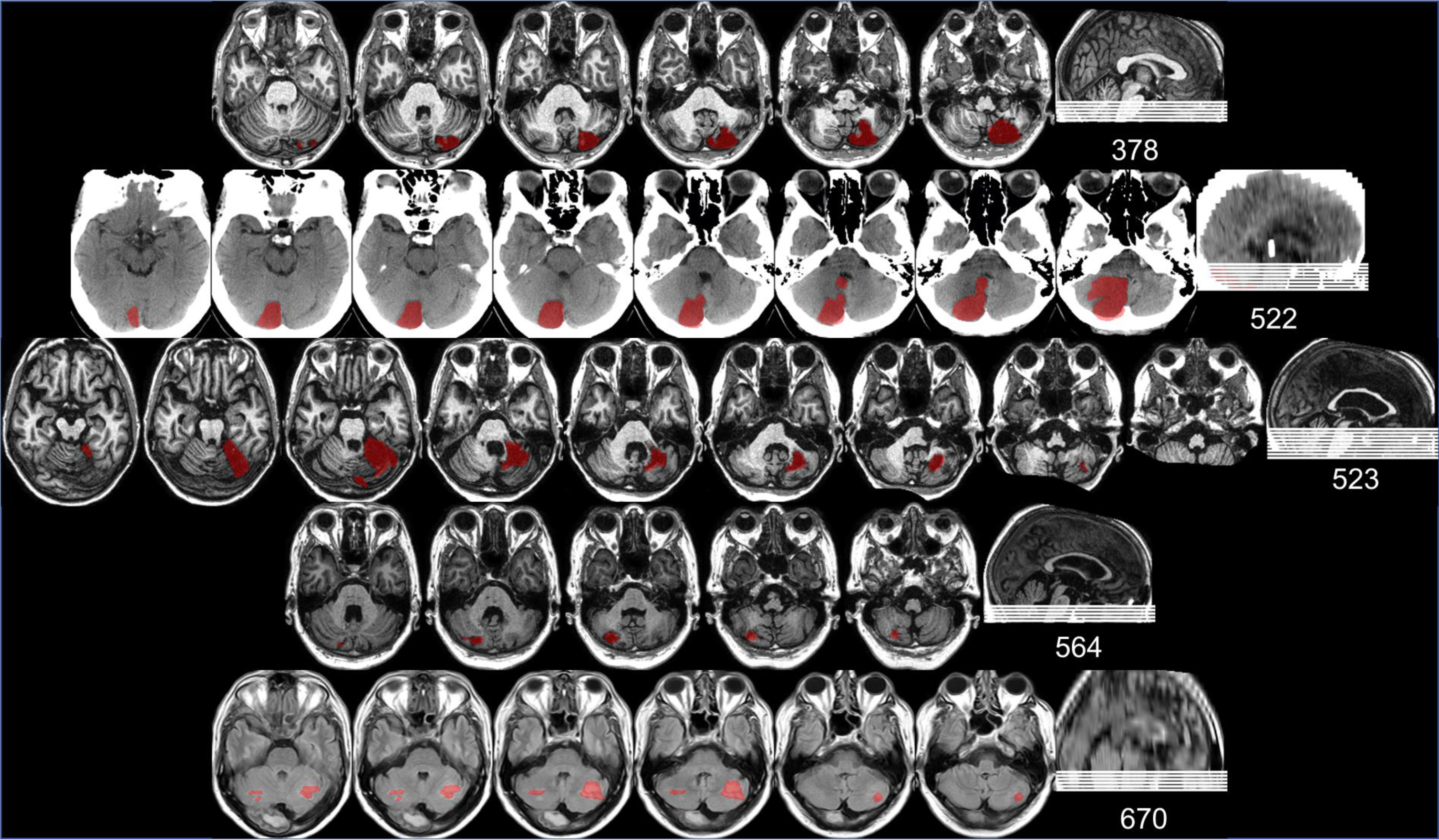
Individual lesion maps for each of the patients (patient number located to the right). See Table 2 in the main manuscript for clinical details for each patient.

**Supplementary Figure 1 Part 3.**
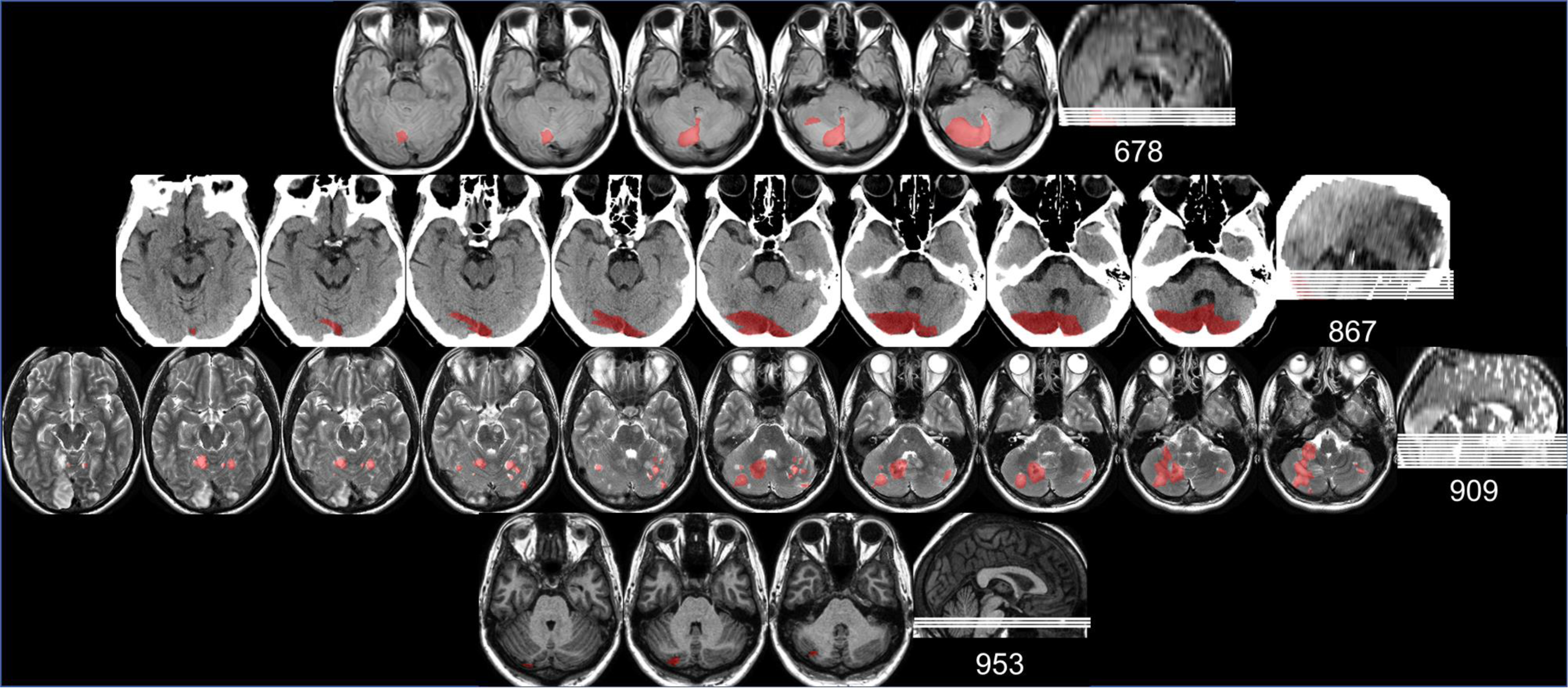
Individual lesion maps for each of the patients (patient number located to the right). See Table 2 in the main manuscript for clinical details for each patient.

**Supplementary Figure 2.**
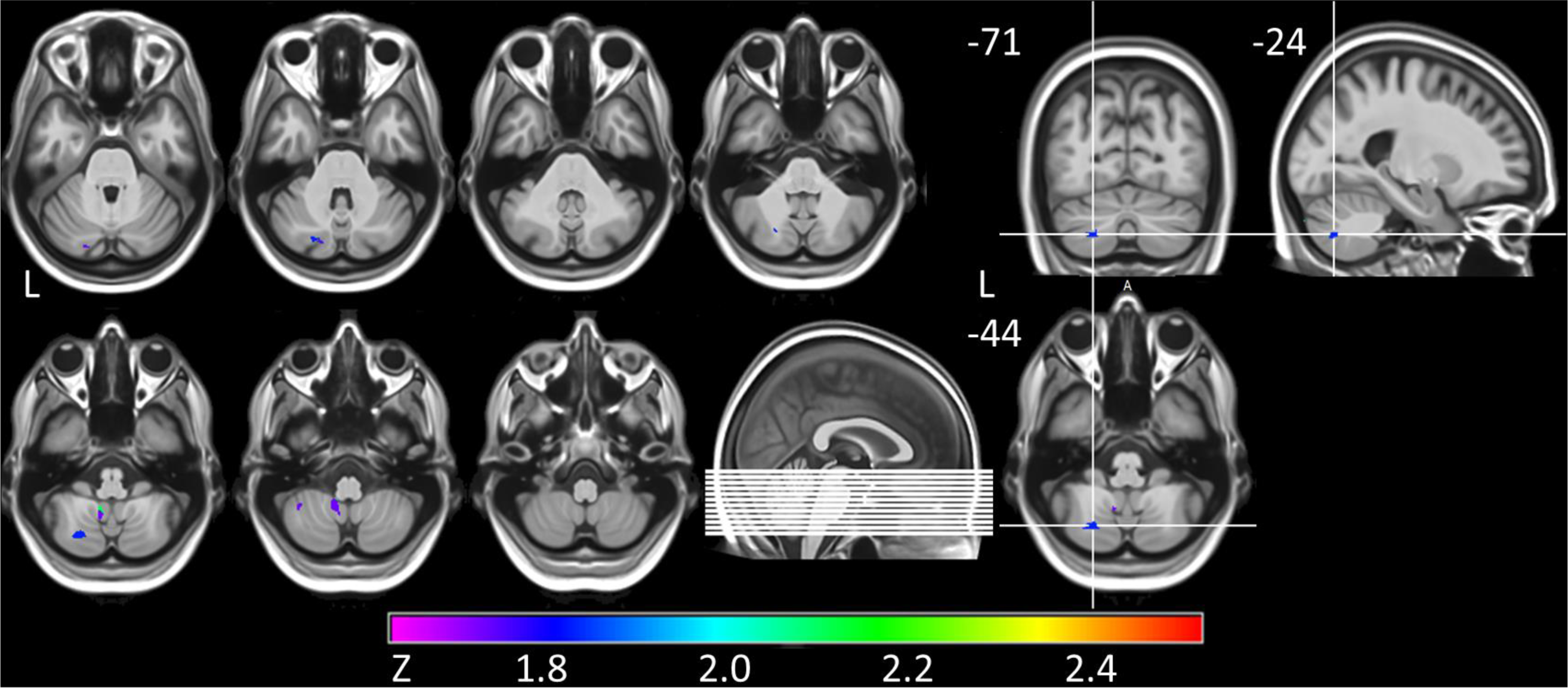
Voxel-based lesion symptom mapping results for the reflexive covert attention task (n=11). The axis indicates the Brunner-Munzel Z-test value (p<.05, uncorrected). The region that is most related to impaired performance is the inferior semilunar lobule (i.e., Crus II) of the left posterior lateral cerebellum.

**Supplementary Figure 3.**
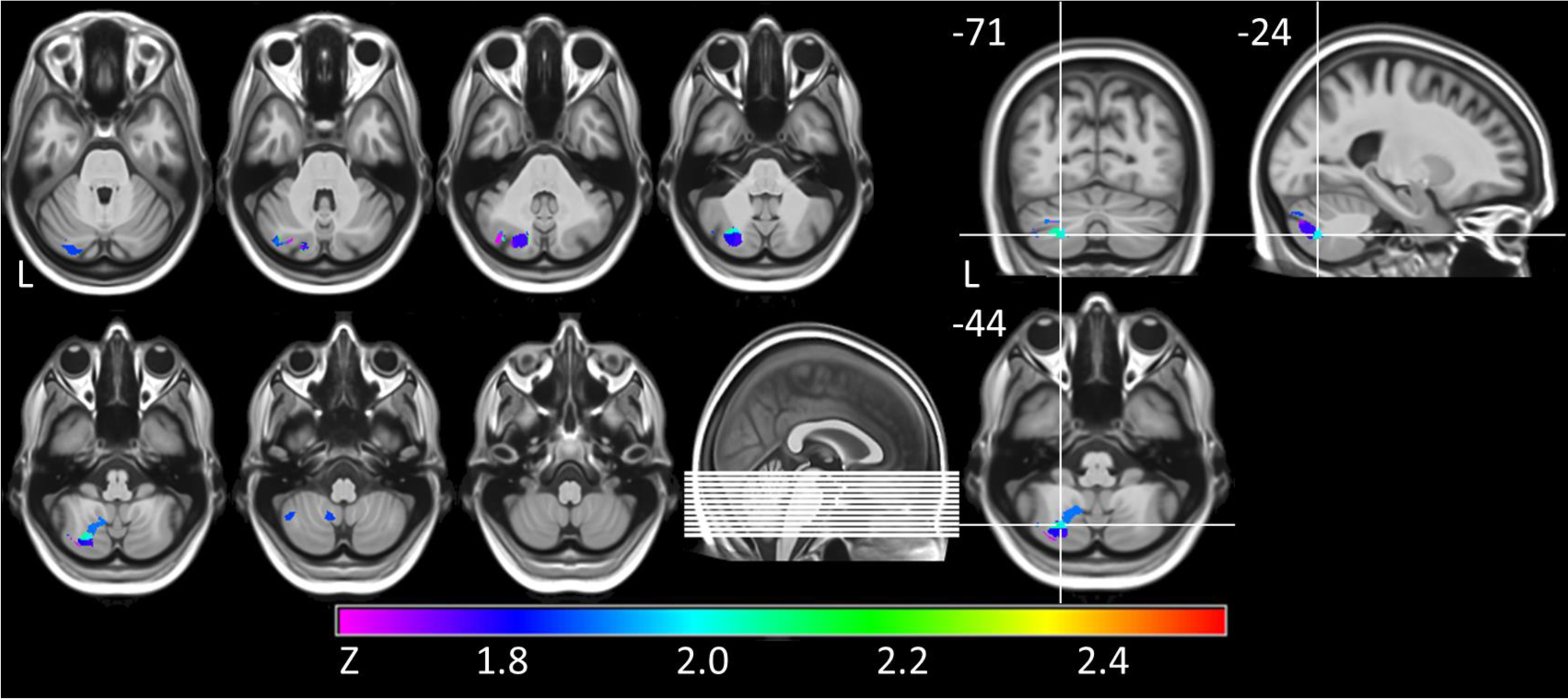
Voxel-based lesion symptom mapping results for the attentional blink (AB) task (n=13). The axis indicates the Brunner-Munzel Z-test value (p<.05, uncorrected). The region that is most related to impaired performance is the inferior semilunar lobule (i.e., Crus II) of the left posterior lateral cerebellum.

